# Downregulation of Semaphorin 4A in keratinocytes reflects the features of non-lesional psoriasis

**DOI:** 10.1101/2024.04.02.587777

**Authors:** Miki Kume, Hanako Koguchi-Yoshioka, Shuichi Nakai, Yutaka Matsumura, Atsushi Tanemura, Kazunori Yokoi, Shoichi Matsuda, Yuumi Nakamura, Naoya Otani, Mifue Taminato, Koichi Tomita, Tateki Kubo, Mari Wataya-Kaneda, Atsushi Kumanogoh, Manabu Fujimoto, Rei Watanabe

## Abstract

Psoriasis is a multifactorial disorder mediated by IL-17-producing T cells, involving immune cells and skin-constituting cells. Semaphorin 4A (Sema4A), an immune semaphorin, is known to take part in T helper type 1/17 differentiation and activation. However, Sema4A is also crucial for maintaining peripheral tissue homeostasis and its involvement in skin remains unknown. Here, we revealed that while Sema4A expression was pronounced in psoriatic blood lymphocytes and monocytes, it was downregulated in the keratinocytes of both psoriatic lesions and non-lesions compared to controls. Imiquimod application induced more severe dermatitis in Sema4A knockout (KO) mice compared to wild-type (WT) mice. The naïve skin of Sema4AKO mice showed increased T cell infiltration and IL-17A expression along with thicker epidermis and distinct cytokeratin expression compared to WT mice, which are hallmarks of psoriatic non-lesions. Analysis of bone marrow chimeric mice suggested that Sema4A expression in keratinocytes plays a regulatory role in imiquimod-induced dermatitis. The epidermis of psoriatic non-lesion and Sema4AKO mice demonstrated mTOR complex 1 upregulation, and the application of mTOR inhibitors reversed the skewed expression of cytokeratins in Sema4AKO mice. Conclusively, Sema4A- mediated signaling cascades can be triggers for psoriasis and targets in the treatment and prevention of psoriasis.

## INTRODUCTION

While the infiltration of immune cells into skin plays a critical role in the development of psoriasis, as evidenced by interleukin (IL) -23/IL-17 axis (Fitch, Harper, Skorcheva, Kurtz, & Blauvelt, 2007; Hawkes, Yan, Chan, & Krueger, 2018; Kim & Krueger, 2017), recent studies have revealed that cells constructing skin structure, such as keratinocytes, fibroblasts, and endothelial cells, also play pivotal roles in the development (Heidenreich, Röcken, & Ghoreschi, 2009; Lowes, Russell, Martin, Towne, & Krueger, 2013; Zhang et al., 2023) and maintenance (Arasa et al., 2019; Francis et al., 2024; Q. Li et al., 2023; Ma et al., 2023; Tan et al., 2015; Zhu et al., 2020) of psoriasis. Among these cells, keratinocytes function as a barrier and produce cytokines, chemokines, and antimicrobial peptides against foreign stimuli, resulting in the activation of immune cells (Ni & Lai, 2020; Zhou, Chen, Cui, Shi, & Guo, 2022).

Semaphorins were initially identified as guidance cues in neural development (Kolodkin, Matthes, & Goodman, 1993; Pasterkamp & Kolodkin, 2003) but are now regarded to play crucial roles in other physiological processes including angiogenesis (Iragavarapu-Charyulu, Wojcikiewicz, & Urdaneta, 2020; Serini, Maione, Giraudo, & Bussolino, 2009), tumor microenvironment (Hui, Tam, Jiao, & Ong, 2019; Jiang et al., 2022; Nakayama, Kusumoto, Nakahara, Fujiwara, & Higashiyama, 2018; Rajabinejad et al., 2020), and immune systems (Garcia, 2019; Kanth, Gairhe, & Torabi-Parizi, 2021; M. Naito & Kumanogoh, 2023). Semaphorin 4A (Sema4A), one of the immune semaphorins, plays both pathogenic and therapeutic roles in autoimmune diseases (Cavalcanti et al., 2020; He et al., 2023), allergic diseases (Maeda et al., 2019), and cancer (Iyer & Chapoval, 2018; Liu et al., 2018; Y. Naito et al., 2023; Pan, Wang, & Ye, 2016). While Sema4A expression on T cells is essential for T helper type 1 differentiation in the murine *Propionibacterium acnes*-induced inflammation model and delayed-type hypersensitivity model (Kumanogoh et al., 2005), Sema4A amplifies only T helper type 17 (Th17)-mediated inflammation in the effector phase of murine experimental autoimmune encephalomyelitis (Koda et al., 2020). Accordingly, in multiple sclerosis, high serum Sema4A levels correlate with the elevated serum IL-17A, earlier disease onset, and increased disease severity (Koda et al., 2020; Nakatsuji et al., 2012). In anti-tumor immunity, research involving human samples and murine models suggests that Sema4A expressed in cancer cells and regulatory T cells promotes tumor progression (Delgoffe et al., 2013; Liu et al., 2018; Pan et al., 2016), while other reports reveal that Sema4A in cancer cells and dendritic cells bolsters anti-tumor immunity by enhancing CD8 T cell activity (Y. Naito et al., 2023; Suga et al., 2021). In addition to these roles in immune reactions, mice with a point mutation in Sema4A develop retinal degeneration, suggesting that Sema4A is also crucial for peripheral tissue homeostasis (Nojima et al., 2013). These reports suggest that the role of Sema4A can differ based on the disease, phase, and involved cells.

Herein, we investigated the roles of Sema4A in the pathogenesis of psoriasis by analyzing skin and blood specimens from psoriatic subjects and using a murine model.

## RESULTS

### Epidermal Sema4A expression is downregulated in psoriasis

The analysis of previously-published single-cell RNA-sequencing data from control (Ctl) and psoriatic lesional (L) skin specimens (Kim et al., 2023) revealed detectable expression of *SEMA4A* in keratinocytes, dendritic cells, and macrophages in both Ctl and L. *SEMA4A* expression was low in neural crest-like cells, fibroblasts, CD4 T cells, CD8 T cells, NK cells, and plasma cells, making the comparison of expression levels impractical (Figure 1A and B; Figure 1-figure supplement 1A-C). Dendritic cells and macrophages showed comparable *SEMA4A* expression levels between Ctl and L (Figure 1C). The adjusted *p*-value (*padj*) for *SEMA4A* in keratinocytes between Ctl and L was 2.83×10^-39^, indicating a statistically significant difference despite not being visually prominent in the volcano plot, which shows comprehensive differential gene expression in keratinocytes (Figure 1C; Figure 1-figure supplement 1D).

**Figure 1:**
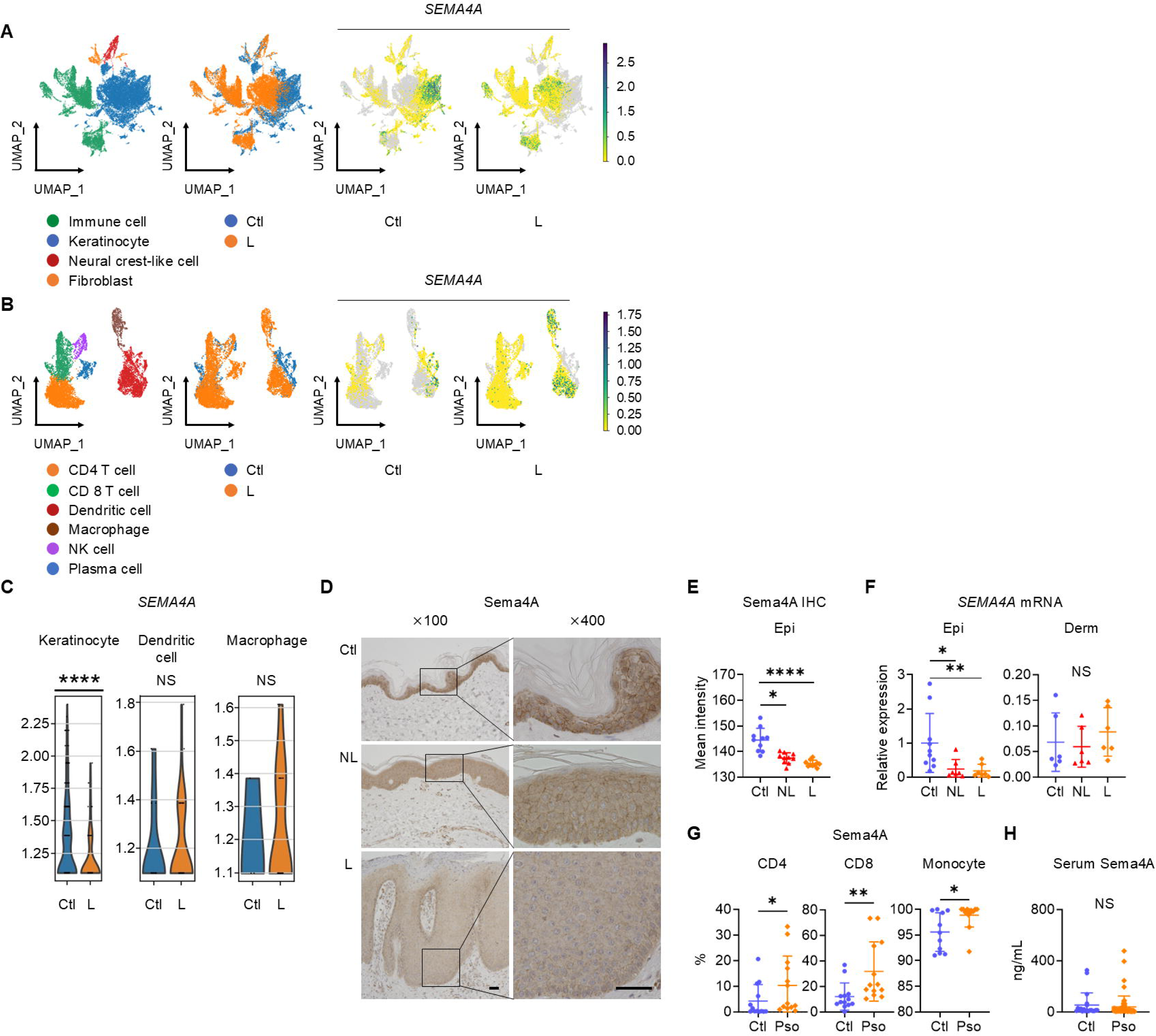
E**p**idermal **Sema4A expression is downregulated in psoriasis.** (**A**) UMAP plots, generated from single-cell RNA-sequencing data (GSE220116), illustrate cell distributions from control (Ctl) and psoriatic lesion (L) samples (*n* = 10 for Ctl, *n* = 11 for L). (**B**) Subclustering of immune cells. (**C**) *SEMA4A* expression in keratinocytes, dendritic cells, and macrophages. \*\*\*\**padj* < 0.001. NS, not significant. Analyzed using Python and cellxgene VIP. (**D**) Representative immunohistochemistry and magnified views showing Sema4A expression in Ctl, psoriatic non-lesion (NL), and L. Scale bar = 50 μm. (**E**) Mean epidermal (Epi) Sema4A intensity in immunohistochemistry (*n* = 10 per group). Each dot represents the average intensity from 5 unit areas per sample. (**F**) Relative *SEMA4A* expression in Epi (*n* = 10 for Ctl, L, *n* = 7 for L and NL) and dermis (Derm, *n* = 6 per group). (**G**) Proportions of Sema4A- expressing cells in blood CD4 T cells (left), CD8 T cells (middle), and monocytes (right) from Ctl and psoriatic (Pso) patients. (*n* = 13 per group in CD4 and CD8, *n* = 11 for Ctl and *n* = 13 for Pso in Monocytes). (**H**) Serum Sema4A levels in Ctl (*n* = 20) and Pso (*n* = 60). **E** to **H**: \**p* < 0.05, \*\**p* < 0.01, \*\*\*\**p* < 0.0001. NS, not significant. Figure 1-source data 1 Excel file containing quantitative data for Figure 1.

Immunohistochemistry of Ctl and psoriasis demonstrated Sema4A expression in keratinocytes (Figure 1D). The staining intensity of Sema4A in epidermis was significantly lower in both non-lesions (NL) and L than in Ctl (Figure 1E). Relative mRNA expression of *SEMA4A* was also decreased in the epidermis of NL and L compared to Ctl, while it remained comparable in the dermis (Figure 1F). In contrast, the proportions of Sema4A-positive cells were significantly higher in blood CD4 and CD8 T cells, and monocytes in psoriasis compared to Ctl (Figure 1G; Figure 1-figure supplement 2). Serum Sema4A levels, measured by enzyme-linked immunosorbent assay (ELISA), were comparable between Ctl and psoriasis (Figure 1H). These findings demonstrate that the expression profile of Sema4A in psoriasis varies across cell types.

### Psoriasis-like dermatitis is augmented in Sema4AKO mice

When psoriasis-like dermatitis was induced in wild-type (WT) mice and Sema4A knockout (KO) mice by imiquimod application on ears (Figure 2A), ear swelling on day 4 was more pronounced in Sema4AKO mice (Figure 2B) with upregulated *Il17a* gene expression (Figure 2C). Flow cytometry analysis of cells isolated from the ears revealed increased proportions of Vγ2^+^ T cells, Vγ2^-^Vγ3^-^ double- negative (DN) γδ T cells, and IL-17A-producing cells of those fractions in Sema4AKO epidermis (Figure 2D and E; Figure 2-figure supplement 1). In Sema4AKO dermis, there was also an increase in the proportions of Vγ2^+^ T cells and IL-17A-producing Vγ2^+^ T cells (Figure 2D and E). These results suggest that Sema4A deficiency in mice accelerates psoriatic profile. IL-17A-producing T cells in skin-draining lymph nodes (dLN) remained comparable between WT mice and Sema4AKO mice (Figure 2F).

**Figure 2:**
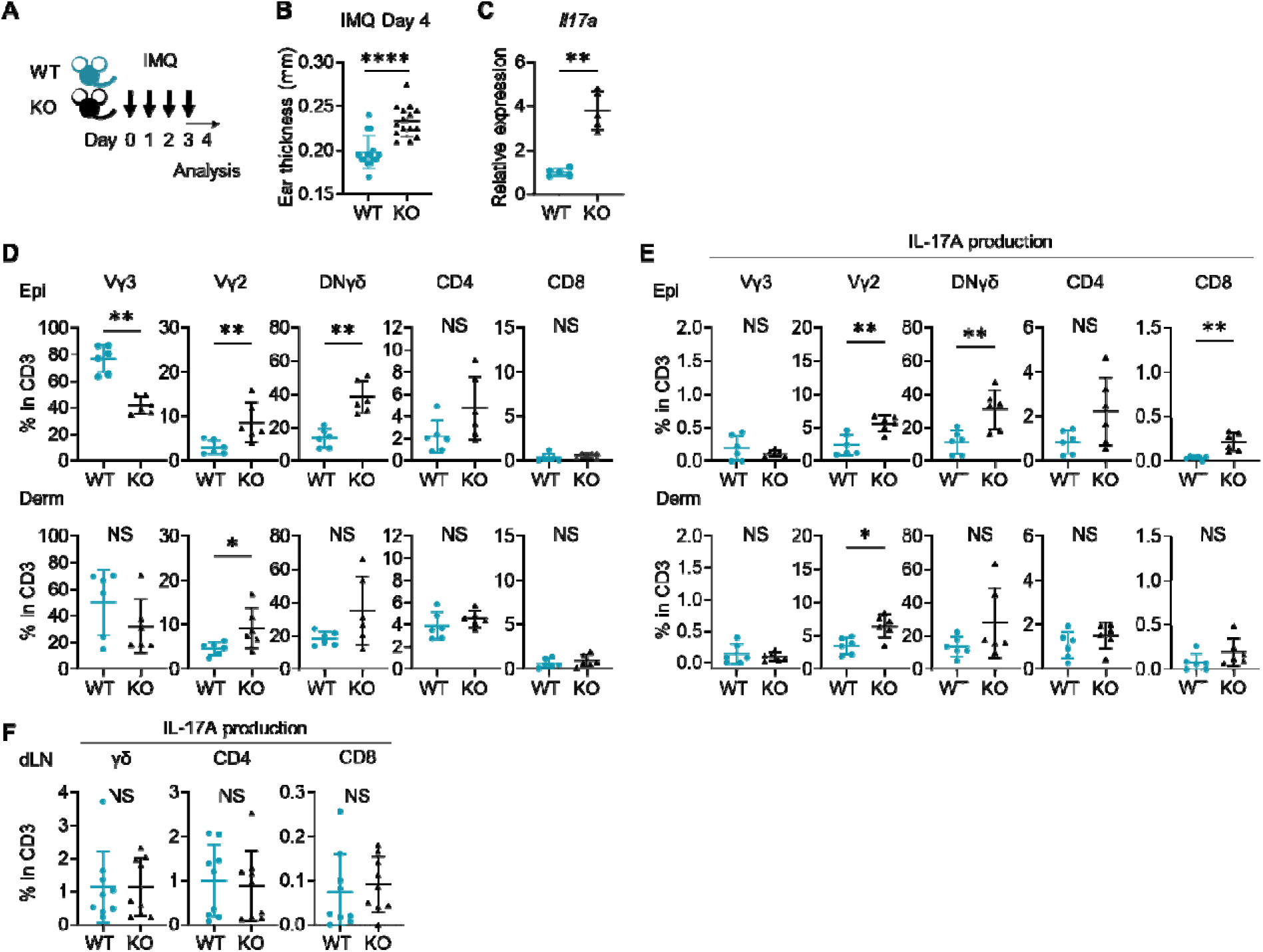
I**m**iquimod**-induced Psoriasis-like dermatitis is augmented in Sema4AKO mice.** (**A**) Experimental scheme. Wild-type (WT, green) mice and Sema4A knockout (KO, black) mice were treated with 10 mg/ear of 5% Imiquimod (IMQ) for 4 consecutive days. Samples for flow cytometry analysis were collected on Day 4. (**B**) Ear thickness of WT mice and KO mice on Day 4 (*n* = 15 per group). (**C**) Relative expression of *Il17a* in epidermis (*n* = 5 per group). (**D** and **E**) The percentages of Vγ3, Vγ2, Vγ2-Vγ3-γδ (DNγδ), CD4, and CD8 T cells (**D**) and those with IL-17A production (**E**) in CD3 fraction in the Epi (top) and Derm (bottom) of WT and KO ears (*n* = 6 per group, each dot represents the average of 4 ear specimens). (**F**) The percentages of IL-17A- producing γδ, CD4, and CD8 T cells in CD3 fraction in skin-draining lymph nodes (dLN) (*n* = 9 per group). **B**-**F**: \**p* < 0.05, \*\**p* < 0.01, \*\*\*\**p* < 0.0001. NS, not significant. Figure 2-source data 1 Excel file containing quantitative data for Figure 2.

Though the imiquimod model is well-established and valuable murine psoriatic model (van der Fits et al., 2009), the vehicle of imiquimod cream can activate skin inflammation that is independent of toll-like receptor 7, such as inflammasome activation, keratinocyte death and interleukin-1 production (Walter et al., 2013). This suggests that the imiquimod model involves complex pathway. Therefore, we subsequently induced IL-23-mediated psoriasis-like dermatitis (Figure2-figure supplement 2A), a much simpler murine psoriatic model, because IL-23 is thought to play a central role in psoriasis pathogenesis (Krueger et al., 2007; Lee et al., 2004). Although ear swelling on day 4 was comparable between WT mice and Sema4AKO mice (Figure2-figure supplement 2B), the epidermis, but not the dermis, was significantly thicker in Sema4AKO mice compared to WT mice (Figure2-figure supplement 2C). We found that the proportion of CD4 T cells among T cells was significantly higher in Sema4A KO mice compared to WT mice, while the proportion of Vγ2 and DNγδ T cells among T cells was comparable between them (Figure 2-figure supplement 2D). On the other hand, focusing on IL-17A-producing cells, the proportion of IL-17A-producing Vγ2 and DNγδ T cells in CD3 fraction in the epidermis was significantly higher in Sema4A KO mice, consistent with the results from imiquimod- induced psoriasis-like dermatitis. (Figure 2-figure supplement 2E).

### Sema4A in keratinocytes may play a role in preventing murine psoriasis-like dermatitis

To investigate the cells responsible for the augmented ear swelling in Sema4AKO mice, bone marrow chimeric mice were next analyzed (Figure 3A). Since it has already been reported that bone marrow cells contain keratinocyte stem cells (Harris et al., 2004; Wu, Zhao, & Tredget, 2010), we confirmed that epidermis of mice deficient in non-hematopoietic Sema4A (WT→KO) showed no obvious detection of *Sema4a*, thereby ruling out the impact of donor-derived keratinocyte stem cells infiltrating the host epidermis (Figure 3-figure supplement 1A). WT→KO mice displayed more pronounced ear swelling than mice with intact Sema4A expression (WT→WT) following imiquimod application (Figure 3B). Similarly, mice with a systemic deficiency of Sema4A (KO→KO) showed severe ear swelling compared to mice deficient in hematopoietic Sema4A (KO→WT) (Figure 3B). Ear swelling was comparable between WT→WT mice and KO→WT mice (Figure 3B). Flow cytometry analysis revealed increased infiltration of IL-17A-producing DNγδ T cells in the epidermis, as well as Vγ2^+^ T cells and IL-17A-producing Vγ2^+^ T cells in the dermis, in WT→KO mice compared to WT→WT mice (Figure 3C; Figure 3-figure supplement 1B). These findings suggest that non-hematopoietic cells, possibly keratinocytes, are primarily responsible for the increased imiquimod-induced Sema4AKO mice ear swelling.

**Figure 3:**
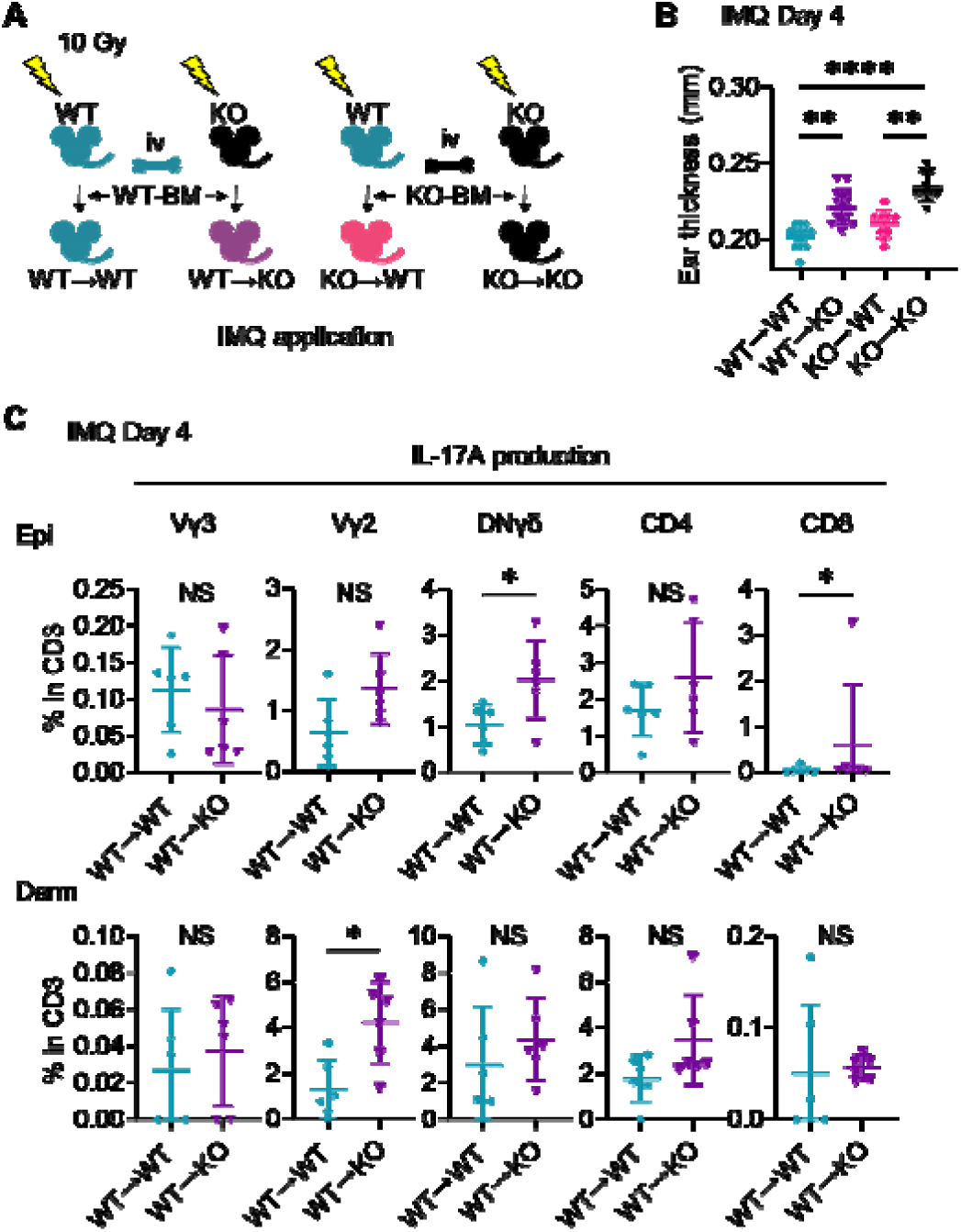
S**e**ma4A **in keratinocytes may play a role in preventing murine psoriasis- like dermatitis.** (**A**) Experimental scheme for establishing BM chimeric mice. (**B**) IMQ Day 4 ear thickness in the mice with the indicated genotypes (*n* = 14 for WT→WT, *n* = 13 for WT → KO, *n* =9 for KO→ WT, *n* = 9 for KO→ KO). (**C**) The percentages of IL-17A- producing Vγ3, Vγ2, DNγδ, CD4, and CD8 T cells in CD3 fraction from IMQ Day 4 Epi (top) and Derm (bottom) of the ears from WT→ WT mice and WT→ KO mice (*n* = 6 per group). Each dot represents the average of 4 ear specimens. **B**-**C**: \**p* < 0.05, \*\**p* < 0.01, \*\*\*\**p* < 0.0001. NS, not significant. Figure 3-source data 1 Excel file containing quantitative data for Figure 3.

### Sema4AKO epidermis is thicker than WT epidermis with increased γδ T17 infiltration

Even without imiquimod application, Sema4AKO ears turned out to be slightly but significantly thicker than WT ears on week 8 while their appearance remained normal (Figure 4A). While epidermal thickness of back skin was comparable at birth (Figure 4B), on week 8, epidermis of Sema4AKO back and ear skin was notably thicker than that of WT mice (Figure 4B), suggesting that acanthosis in Sema4AKO mice is accentuated post-birth. Dermal thickness remained comparable between WT mice and Sema4AKO mice at both times (Figure 4B). The epidermis of WT ear at week 8 showed significantly higher *Sema4a* mRNA expression compared to the dermis (Figure 4C). Based on these observations, Sema4A appears to play a more pronounced role in epidermis than in dermis.

**Figure 4:**
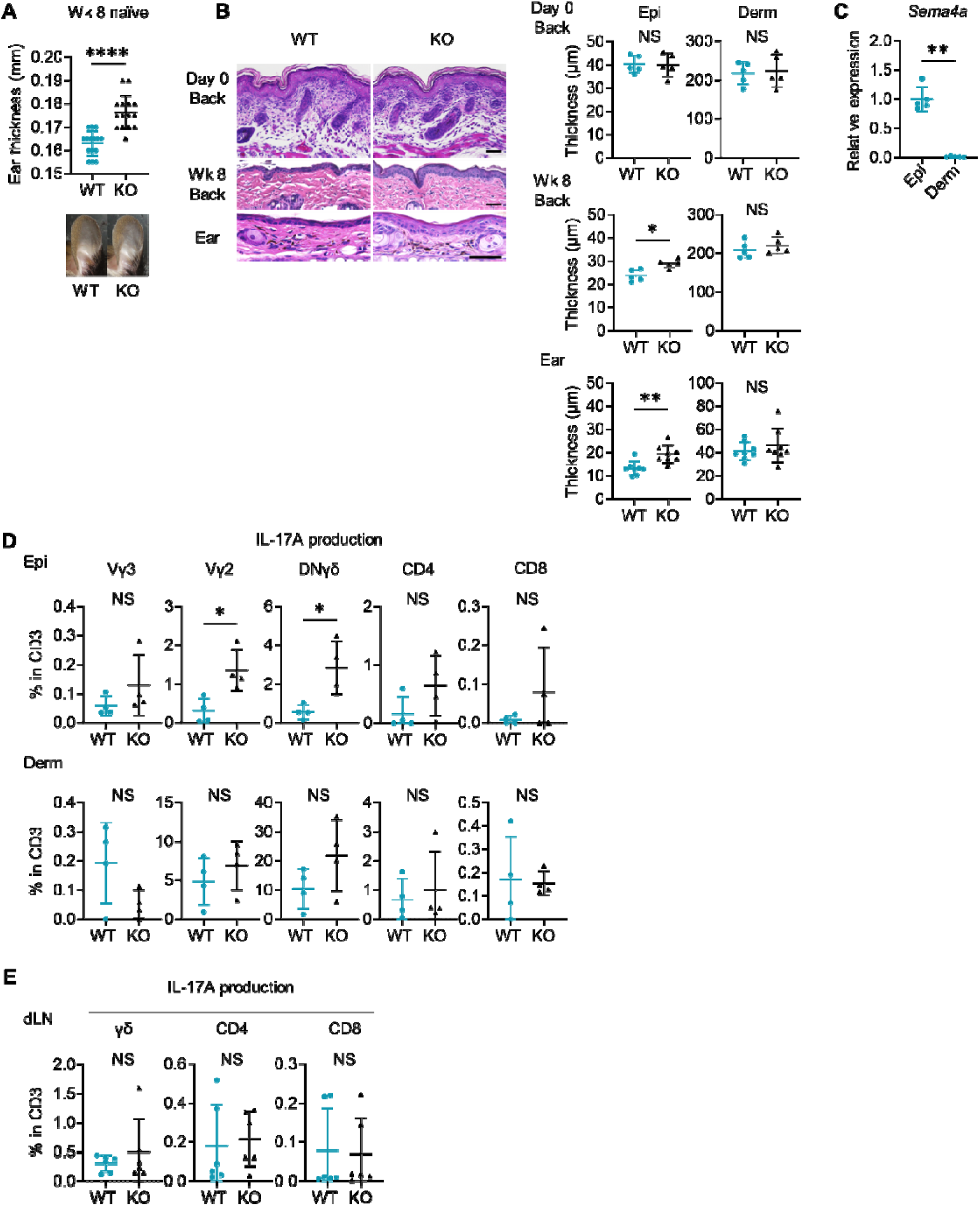
N**a**ïve **Sema4AKO epidermis is thicker than WT epidermis with increased** γδ **T17 infiltration.** (**A**) Ear thickness of WT mice and KO mice at week (Wk) 8 (*n* = 15 per group) and representative images. (**B**) Left: representative Hematoxylin and eosin staining of Day 0 back and Wk 8 back and ear. Scale bar = 50 μm. Right: Epi and Derm thickness in Day 0 back (*n* = 5) and Wk 8 back (*n* = 5) and ear (*n* = 8). (**C**) Relative *Sema4a* expression in WT Epi and Derm (*n* = 5 per group). (**D**) The percentages of the IL-17A-producing Vγ3, Vγ2, DNγδ, CD4, and CD8 T cells in CD3 fraction (*n* = 4 per group) in Epi (top) and Derm (bottom). Each dot represents the average of 4 ear specimens. (**E**) The graphs showing the percentages IL-17A-producing γδ, CD4, and CD8 T cells in CD3 fraction from draining LN (dLN) of WT mice and Sema4AKO mice (*n* = 6 per group). **A**-**E**: \**p* < 0.05, \*\**p* < 0.01, \*\*\*\**p* < 0.0001. NS, not significant. Figure 4-source data 1 Excel file containing quantitative data for Figure 4.

Sema4AKO epidermis exhibited increased expression of *Ccl20, Tnf*α, and *Il17a* and a trend of upregulation of *S100a8* compared to WT epidermis (Figure 4-figure supplement 1A). These differences were not observed in dermis (Figure 4-figure supplement 1A). Flow cytometry analysis revealed increased infiltration of γδ T cells in Sema4AKO ear (Figure 4-figure supplement 1B). These cells predominantly expressed resident memory T cell (T_RM_)-characteristic molecules, CD69 and CD103 (Figure 4-figure supplement 1B). Sema4AKO skin also had a higher number of T_RM_ in both CD4 and CD8 T cells (Figure 4-figure supplement 1B). The percentages of Vγ2^+^ T cells, DNγδ T cells, and CD8 T cells in epidermis were higher in Sema4AKO mice than in WT mice, which was not the case in dermis (Figure 4-figure supplement 2A). The proportion of Vγ3^+^ dendritic epidermal T cells was comparable between WT mice and Sema4AKO mice (Figure 4-figure supplement 2A). Epidermal Vγ2^+^ T cells and DNγδ T cells in Sema4AKO mice showed higher IL-17A-producing capability (Figure 4D), while IFNγ and IL-4 production was comparable between WT mice and Sema4AKO mice (Figure 4-figure supplement 2B and C). Conversely, the frequency of IL-17A- producing T cells from dLN was comparable (Figure 4E). The production of IL-17A and IFNγ from splenic T cells under T17-polarizing conditions remained consistent between WT mice and Sema4AKO mice (Figure 4-figure supplement 3).

Taken together, it is suggested that T17 cells are specifically upregulated in epidermis, indicating that the epidermal microenvironment plays a pivotal role in facilitating the increased T cell infiltration observed in naïve Sema4AKO mice.

### Sema4AKO skin shares features with human psoriatic NL

Previous literatures have identified certain features common to psoriatic L and NL, such as thickened epidermis (Figure 5-figure supplement 1) (Gallais Sérézal et al., 2019), CCL20 upregulation (Gallais Sérézal et al., 2019), and accumulation of IL-17A- producing T cells (Vo et al., 2019), which were detected in Sema4AKO mice.

**Figure 5:**
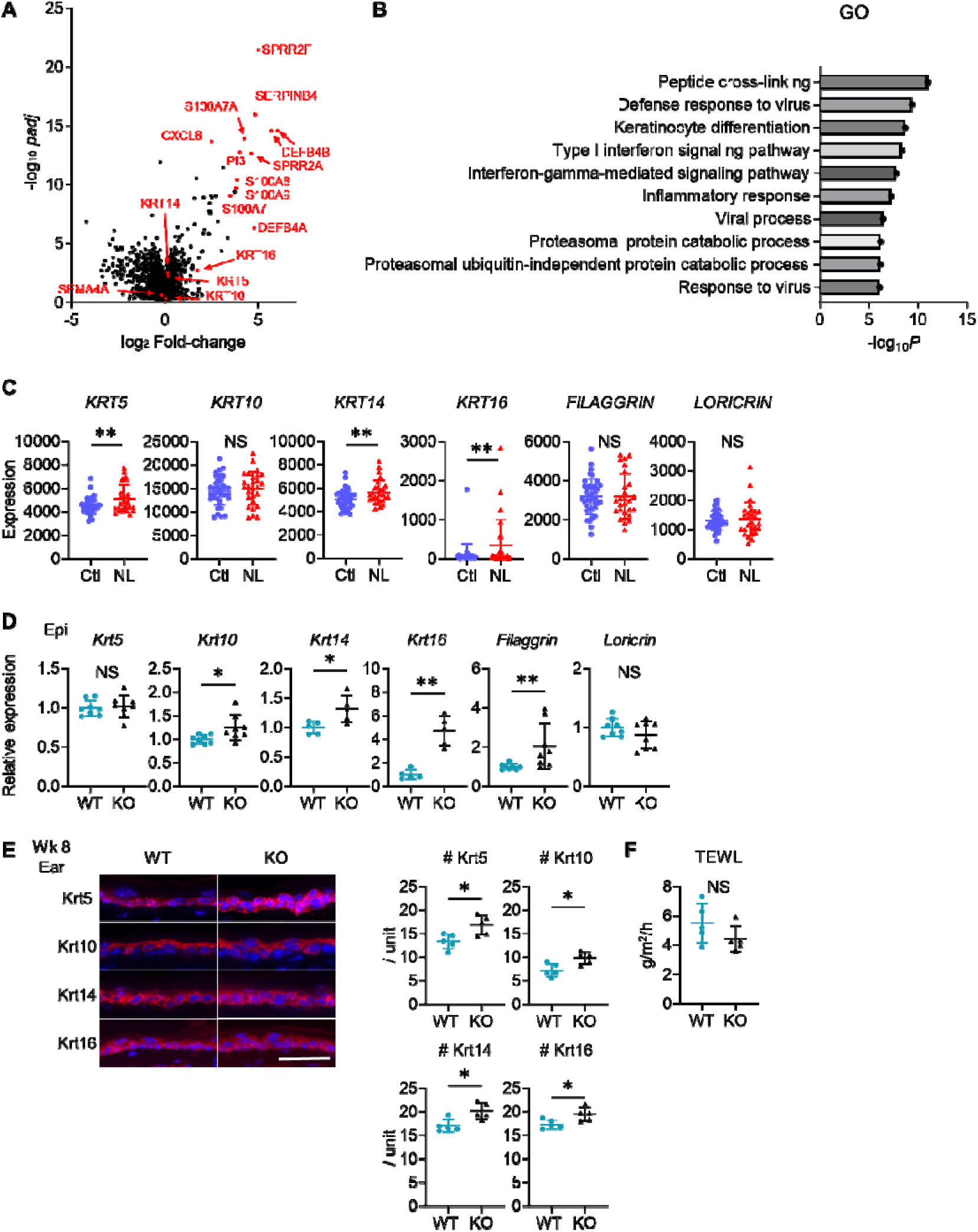
S**e**ma4AKO **skin shares the features of human psoriatic NL.** (**A** and **B**) The volcano plot (**A**) and Gene ontology (GO) analysis (**B**), generated from RNA-sequencing data (GSE121212) using RaNA-seq, display changes in gene expression in psoriatic NL compared to Ctl. (**C**) The difference in the expression of epidermal differentiation markers between Ctl and NL (*n* = 38 for Ctl, *n* = 27 for NL) was calculated with the transcripts per million values. \*\**padj* < 0.01. NS, not significant. (**D**) Relative gene expression of epidermal differentiation markers between wk 8 Epi of WT mice and KO mice (*n* = 5 for *Krt14* and *Krt16*, *n* = 8 for *Krt5*, *Krt10*, *Filaggrin*, and *Loricrin*). (**E**) Left: Representative immunofluorescence pictures of Krt5, Krt10, Krt14, and Krt16 (red) overlapped with DAPI. Scale bar = 50 μm. Right: Accumulated graphs showing the numbers of Krt5, Krt10, Krt14, and Krt16 positive cells per 100 μm width (*n* = 5 per group) of wk 8 ear (right). Each dot represents the average from 5 unit areas per sample. (**F**) Transepidermal water loss (TEWL) in back skin of WT mice and KO mice at wk 8 (*n* = 5 per group). **D**-**F**: \**p* < 0.05, \*\**p* < 0.01. NS, not significant. Figure 5-source data 1 Excel file containing quantitative data for Figure 5.

Gene expression analysis using public RNA-sequencing data (Tsoi et al., 2019) with RaNA-seq (Prieto & Barrios, 2019) showed upregulation of keratinization and antimicrobial peptide genes in NL compared to Ctl (Figure 5A). Gene Ontology analysis highlighted an upregulation in biological processes, predominantly in peptide cross- linking involved in epidermis formation and keratinocyte differentiation, with a secondary increase in the defense response to viruses in NL (Figure 5B). While the expression of *Keratin (KRT) 10* was comparable between NL and Ctl, upregulation in *KRT5*, *KRT14*, and *KRT16* was observed in NL (Figure 5C).

In the murine model, relative expression levels of *Krt10*, *Krt14*, *Krt16*, and *Filaggrin* were elevated in Sema4AKO epidermis. (Figure 5D). Immunofluorescence analysis showed that Sema4AKO epidermis had a higher density of keratinocytes positive for Krt5, Krt10, Krt14, and Krt16 compared to WT epidermis (Figure 5E). This upregulation was not observed in back skin at birth (Figure 5-figure supplement 2). Comparable transepidermal water loss between WT mice and Sema4AKO mice indicated preserved skin barrier function in Sema4AKO mice (Figure 5F).

Based on these results, it is implied that the epidermis of human psoriatic NL and Sema4AKO mice exhibit shared pathways, potentially leading to an acanthotic state. Combined with the observed acanthosis and increased T17 infiltration in Sema4AKO mice, Sema4AKO skin is regarded to demonstrate the features characteristic of human psoriatic NL.

### mTOR signaling is upregulated in the epidermis of psoriatic NL and Sema4AKO mice

Previous reports have shown that mTOR pathway plays a critical role in maintaining epidermal homeostasis, as evidenced by mice with keratinocyte-specific deficiencies in Mtor, Raptor, or Rictor, which exhibit a hypoplastic epidermis with impaired differentiation and barrier formation (Asrani et al., 2017; Ding et al., 2016; Ding et al., 2020). We thus investigated mTOR pathway in both human and murine epidermis.

Immunohistochemical analyses highlighted the increase in phospho-S6 (p-S6), indicating the upregulation of mTOR complex (C) 1 signaling, in the epidermal upper layers of both L and NL compared to Ctl (Figure 6A). The activation of mTORC2 in the epidermis, inferred from phospho-Akt (p-Akt), was scarcely detectable in L, NL, and Ctl (Figure 6A). In the epidermis of WT mice and Sema4AKO mice, the upregulation of both mTORC1 and mTORC2 signaling was observed in Sema4AKO epidermis by immunohistochemistry (Figure 6B). Western blot analysis showed upregulated mTORC1 signaling in Sema4AKO epidermis compared to WT epidermis (Figure 6C). The enhancement of mTORC1 and mTORC2 signaling became obvious in the Sema4AKO epidermis after developing psoriatic dermatitis by imiquimod application (Figure 6D).

**Figure 6:**
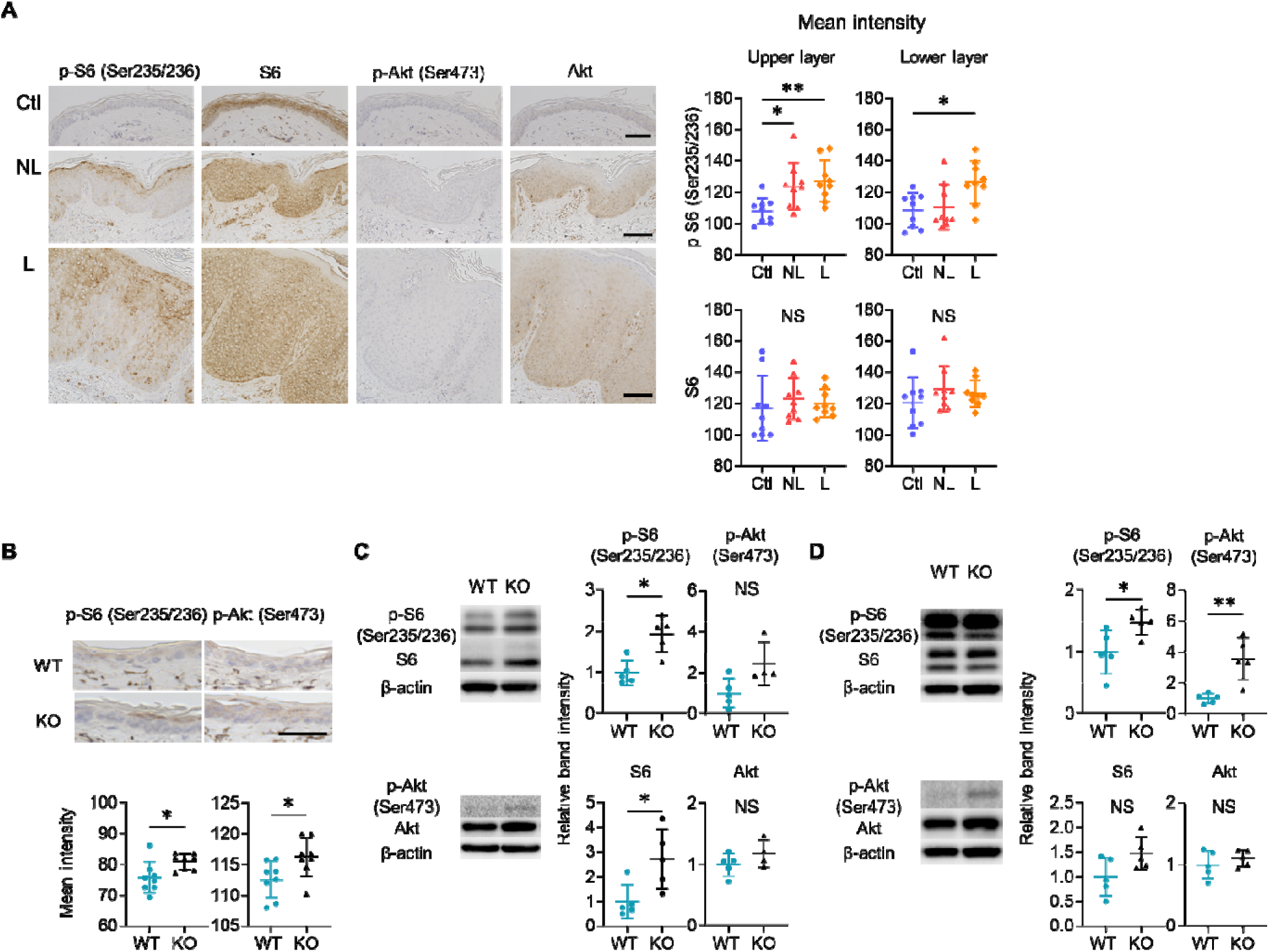
mTOR signaling is upregulated in the epidermis of psoriatic NL and Sema4AKO mice. (A) Representative results of immunohistochemistry displaying cells positive for p-S6 (Ser235/236), S6, p-Akt (Ser473), and Akt in Ctl, NL, and L. The graphs of accumulated data show the mean intensity of p-S6 and S6 in the upper and lower epidermal layers (*n* = 9 per group). Scale bar = 100 μm. Each dot represents the average mean intensity from 5 unit areas per sample. (B) The mean intensity of p-S6 (Ser235/236) and p-Akt (Ser473), detected by immunohistochemistry in the epidermis of WT mice and KO mice, were analyzed. Scale bar = 50 μm. Each dot represents the average intensity from 5 unit areas per sample (*n* = 8 per group). (C and D) Immunoblotting of p-S6 (Ser235/236), S6, p-Akt (Ser473), and Akt in tissue lysates from epidermis without treatment (C) and with IMQ treatment for consecutive 4 days (D) (*n* = 5 per group, except for p-Akt and Akt in C, for which *n* = 4). A-D: \**p* < 0.05, \*\**p* < 0.01. NS, not significant. Figure 6-source data 1 Excel file containing quantitative data for Figure 6.

### Inhibition of mTOR signaling modulates cytokeratin expression in Sema4AKO mice

To investigate the contribution of mTORC1 and mTORC2 signaling in the development of psoriatic features in the Sema4AKO epidermis, mTORC1 inhibitor rapamycin and mTORC2 inhibitor JR-AB2-011 were intraperitoneally applied to Sema4AKO mice for 14 days. Although epidermal thickness remained unchanged by the inhibitors (Figure 7A and B), relative gene expression of *Krt5* was significantly upregulated and that of *Krt16* was significantly downregulated after rapamycin application (Figure 7C). While the upregulation of *Il17a* in Sema4AKO epidermis was not clearly modified by rapamycin (Figure 7C), immunofluorescence revealed the decrease in the number of CD3 T cells in Sema4AKO epidermis by rapamycin (Figure 7D). We additionally conducted topical application of rapamycin gel and vehicle gel on the left and right ears of Sema4AKO mice, respectively. Although there were no detectable changes in epidermal thickness and epidermal T cell counts, the upregulation of *Krt5* and downregulation of *Krt16* was observed again (Figure 7-figure supplement 1). Conversely, the application of JR-AB2-011 resulted in decreased expression of *Krt5*, *Krt10*, and *Krt14* with a trend towards increased *Krt16* expression (Figure 7E). JR- AB2-011 did not influence the number of infiltrating T cells in the epidermis (Figure 7F). Next, we investigated whether intraperitoneal rapamycin treatment effectively downregulates inflammation in the IMQ-induced murine model of psoriasis in Sema4AKO mice (Figure 7-figure supplement 2A). Rapamycin significantly reduced epidermal thickness compared to vehicle treatment (Figure 7-figure supplement 2B). Additionally, rapamycin treatment downregulated the expression of *Krt10*, *Krt14*, and *Krt16* (Figure 7-figure supplement 2C). While the upregulation of *Il17a* in the Sema4AKO epidermis in IMQ model was not clearly modified by rapamycin (Figure 7-figure supplement 2C), immunofluorescence revealed a decrease in the number of CD3 T cells in Sema4AKO epidermis by rapamycin (Figure 7-figure supplement 2D). In the naive states, mTORC1 primarily regulates keratinocyte proliferation, whereas mTORC2 mainly involved in the keratinocyte differentiation through Sema4A-related signaling pathways. Conversely, in the psoriatic dermatitis state, rapamycin downregulated both keratinocyte differentiation and proliferation markers. The observed similarities in *Il17a* expression following treatment with rapamycin and JR-AB2-011, regardless of additional IMQ treatment, suggest that *Il17a* production is not significantly dependent on Sema4A-related mTOR signaling.

**Figure 7:**
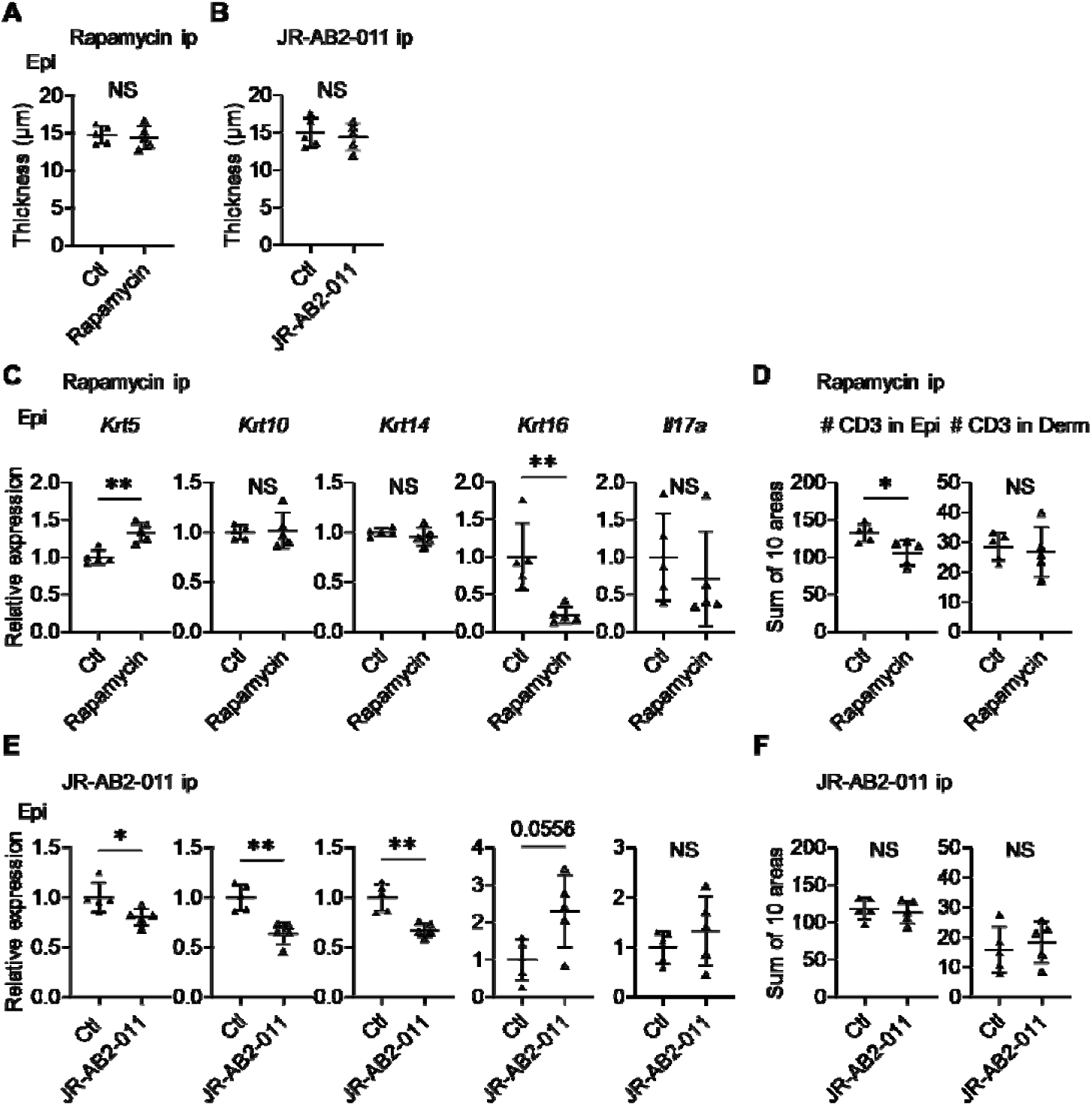
Inhibitors of mTOR signaling modulate the expression of cytokeratins in Sema4AKO mice. (A and B) Epidermal thickness of Sema4AKO mice treated intraperitoneally with vehicle (Ctl) or rapamycin (A), and Ctl or JR-AB2-011 (B) (*n* = 5 per group). (C and D) Relative expression of keratinocyte differentiation markers and *Il17a* in Sema4AKO Epi (C), and the number of T cells in Epi and Derm under Ctl or rapamycin (D) (*n* = 5 per group). (E and F) Relative expression of keratinocyte differentiation markers and *Il17a* in Sema4AKO Epi (E), and the number of T cells in Epi and Derm under Ctl or JR- AB2-011 (F) (*n* = 5 per group). D and F: Each dot represents the sum of numbers from 10 unit areas across 3 specimens. A-F: \**p* < 0.05, \*\**p* < 0.01. NS, not significant. Figure 7-source data 1 Excel file containing quantitative data for Figure 7.

## DISCUSSION

Recent studies have highlighted the significance of IL-17A-producing T_RM_ in psoriatic NL (Cheuk et al., 2014; Gallais Sérézal et al., 2018; Vo et al., 2019; Vu, Koguchi-Yoshioka, & Watanabe, 2021). This condition is also characterized by epidermal thickening with elevated CCL20 expression (Gallais Sérézal et al., 2019) and a distinct cytokeratin expression pattern, marked by increased levels of Krt5, Krt14, and Krt16 (Tsoi et al., 2019) (Figure 5C). Here, we identified that murine Sema4A deficiency could induce these features of psoriatic NL.

Sema4A expression in epidermis, but not in dermis, was diminished in psoriatic L and NL compared to Ctl. While Sema4A expression on blood immune cells was upregulated in psoriasis, its serum levels were comparable between Ctl and psoriasis. Despite the reported involvement of increased Sema4A expression on immune cells in the pathogenesis of multiple sclerosis (Koda et al., 2020; Nakatsuji et al., 2012), our results suggest that the diminished expression of Sema4A in skin-constructing cells plays a more prominent role in the pathogenesis of psoriasis than its increased expression on immune cells.

In our murine model, regardless of imiquimod application, IL-17A-producing T cells were increased in Sema4AKO skin although their frequency was comparable in dLN of WT mice and Sema4AKO mice. Additionally, the potential for *in vitro* T17 differentiation did not differ between T cells from WT mice and Sema4AKO mice. These findings suggest that the absence of Sema4A in the skin microenvironment plays a crucial role in the localized upregulation of IL-17A-producing T cells in Sema4AKO mice.

It is well-documented that upregulated mTORC1 signaling promotes the pathogenesis of psoriasis (Buerger, 2018; Karagianni, Pavlidis, Malakou, Piperi, & Papadavid, 2022; Ruf, Andreoli, Itin, Pluschke, & Schmid, 2014), and that rapamycin can partially ameliorate the disease activity in psoriatic subjects and murine models (Bürger et al., 2017; Gao & Si, 2018; Reitamo et al., 2001). Our results using Sema4AKO mice revealed that inhibition of mTORC1 leads to the downregulation of *Krt16*, and inhibition of mTORC2 leads to the downregulation of undifferentiated keratinocytes, while the Sema4AKO epidermal thickening and the upregulated IL-17A signaling are not reversed by mTOR blockade in the naïve state. It is plausible that the downregulation of Sema4A can lead to the upregulation of mTORC1 and mTORC2 signaling in keratinocytes, and the augmented signaling leads to the psoriatic profile of proliferation and differentiation of keratinocytes, which is part of the psoriatic NL disposition.

This study has limitations. Sema4A expression in skin cells other than keratinocytes was not thoroughly investigated. However, since single-cell RNA- sequencing has shown that Sema4A is predominantly expressed in keratinocytes, dendritic cells, and macrophages, with a notable reduction in keratinocytes in psoriasis, we infer that keratinocytes are the primary cells responsible for the psoriatic features resulting from Sema4A downregulation. We were not able to determine whether Sema4A functions as a ligand or a receptor in epidermis (Ito & Kumanogoh, 2016; Kumanogoh et al., 2002; Lu et al., 2018; Sun et al., 2017) in this study, either. We were not able to reveal how Sema4A expression is regulated. Although we showed that downregulation of Sema4A is related to the abnormal cytokeratin expression observed in psoriasis, we could not determine the relationships between Sema4A expression and the essential molecules upregulated in psoriatic keratinocytes. While both mTORC1 and mTORC2 signals are upregulated in Sema4AKO epidermis, we were not able to confirm mTORC2 signaling from human skin due to technical limitation and sample limitation, while the results from augmented mTORC2 signal in Sema4AKO mice and the normalization of *Krt5* and *Krt14* by mTORC2 inhibitor imply the involvement of mTORC2 signal in psoriatic epidermis. The role of Sema4A other than mTOR signaling, which may be involved in the regulation of T17 induction in skin, is not discussed, either. These limitations should be overcome in the near future.

In summary, epidermal Sema4A downregulation can reflect the psoriatic non- lesional features. Thus, targeting the downregulated Sema4A and upregulated mTOR signaling in psoriatic epidermis can be a promising strategy for psoriasis therapy and modification of psoriatic diathesis in NL for the prevention of disease development.

## MATERIALS AND METHODS

### Human sample collection

Psoriatic L and NL skin specimens were acquired from 17 psoriasis patients. Ctl specimens were obtained from 19 subjects who underwent tumor resection or reconstructive surgery. For epidermal-dermal separation, specimens were incubated overnight at 4℃ with 2.5 mg/mL dispase Ⅱ (Wako, Osaka, Japan) in IMDM (Wako). Blood samples were collected from 73 psoriasis and 33 Ctl. In addition to ELISA (Nakatsuji et al., 2012), peripheral blood mononuclear cells were isolated using Ficoll- Paque PLUS density gradient media (Cytiva, Tokyo, Japan). Patient details are provided in Supplemental Table 1.

### Mice

C57BL/6J WT mice were procured from CLEA Japan (Tokyo, Japan). Sema4AKO mice with C57BL/6J background were generated as previously described (Kumanogoh et al., 2005). We examined female mice in order to reduce the result variation. Neonatal mice and female mice aged 8 to 12 weeks, maintained under specific pathogen-free conditions, were used.

To develop psoriasis-like dermatitis, 10 mg of 5% imiquimod cream (Mochida, Tokyo, Japan) was applied to both ears for 4 consecutive days. To induce IL-23- mediated psoriasis-like dermatitis, 20 μl of phosphate-buffered saline containing 500 ng of recombinant mouse IL-23 (BioLegend, San Diego, CA) was injected intradermally into both ears of anesthetized mice using a 29-gauge needle for 4 consecutive days. Ear thickness was measured using a thickness gauge (Peacock, Tokyo, Japan).

For bone marrow reconstitution, 2 × 10^5^ bone marrow cells obtained from the indicated strains were intravenously transferred to recipient mice that had received a single 10 Gy irradiation. The reconstituted mice were subjected to the experiments in 8 weeks.

In the specified experiments, skin specimens were separated into epidermis and dermis by 5 mg/mL dispase (Wako) in IMDM for 30 minutes at 37℃.

Transepidermal water loss measurements were performed on the back skin of mice at week 8 using a Tewameter (Courage and Khazaka Electronic GmbH, Cologne, Germany), according to the manufacturer’s instructions.

### Immunofluorescence and immunohistochemistry

Specimens were fixed in 4% paraformaldehyde phosphate buffer solution (Wako), embedded in paraffin, and sliced into 3-μm thickness on glass slides. After deparaffinization and rehydration, antigen retrieval was performed using citrate buffer (pH 6.0, Nacalai tesque, Kyoto, Japan) or TE buffer (pH 9.0, Nacalai tesque).

For immunohistochemistry, samples were incubated with 3% H_2_O_2_ (Wako) for 5 minutes. After blocking (Agilent, Santa Clara, CA), the specimens were incubated with the indicated primary antibodies under specified conditions (Supplemental Table 2). The specimens were applied with the Dako REAL EnVision Detection System, Peroxidase/DAB, Rabbit/Mouse, HRP kit (Agilent) and counterstained with hematoxylin (Wako). For immunofluorescence, the blocked specimens were incubated with the indicated primary antibodies followed by the secondary antibodies (Table E4). Mounting medium with DAPI (Vector Laboratories, Burlingame, CA) was used. Slides were observed using a fluorescence microscope (BZ-X700, Keyence, Osaka, Japan). The staining intensity was measured using Fiji software (ImageJ, National Institutes of Health, Bethesda, MD) over lengths of 200 μm in human samples and 100 μm in murine samples. This measurement was taken from five areas of each specimen, and the average score is presented. The average thickness of epidermis and dermis from 10 spots is presented. The number of epidermal cells positive for the indicated cytokeratins was counted per 100 μm width, with the average number from 5 areas being presented. Additionally, the numbers of CD3-positive cells in murine ears were counted across 10 fields of 400 μm width, with the total sum being presented.

### Quantitative reverse transcription polymerase chain reaction (qRT-PCR)

Total RNA was extracted from homogenized skin tissue using Direct-zol RNA MiniPrep Kit (ZYMO Research, Irvine, CA). cDNA was synthesized by High-Capacity RNA-to-cDNA^TM^ Kit (Thermo Fisher Scientific, Waltham, MA). qPCR was performed using TB Green Premix Ex Taq^TM^ II (Takara-bio, Shiga, Japna) on a ViiA 7 Real-Time PCR System (Thermo Fisher Scientific). The primers are listed in Supplemental Table 3. All samples were run in triplicate, and the median CT value was calculated. Relative gene expression levels were normalized to the housekeeping gene *GAPDH* using the ΔΔCT technique.

### Flow cytometry analysis

Single-cell suspensions from murine skin-dLN and spleen were prepared by grinding, filtering, and lysing red blood cells (BioLegend). Skin specimens were minced and digested with 3 mg/mL collagenase type III (Worthington Biochemical Corporation, Lakewood, NJ) in RPMI 1640 medium (Wako) at 37℃ for 10 minutes for epidermis, and 30 minutes for dermis or whole skin. Cells were surface-stained with directly conjugated monoclonal antibodies (Supplemental Table 4). Dead cells were identified using LIVE/DEAD^TM^ Fixable Dead Cell Stain Kit (Thermo Fisher Scientific). To evaluate cytokine production, cells were stimulated with Phorbol 12-Myristate 13- Acetate (PMA; 50 ng/mL, Wako) and ionomycin (1000 ng/mL, Wako), plus BD Golgiplug^TM^ (BD Biosciences, San Jose, CA) for 4 hours before surface staining. Fixation, permeabilization, and intracellular cytokine staining were performed using BD Cytofix/Cytoperm^TM^ Fixation/Permeabilization Kit (BD Biosciences) according to the manufacturer’s protocol. In specified experiments, the numbers of each cell subset per ear were estimated using CountBright^TM^ Absolute Counting Beads (Thermo Fisher Scientific). Sample analysis was conducted using BD FACSCanto II (BD Biosciences), and data were analyzed with Kaluza software (Beckman Coulter, Brea, CA).

### *In vitro* Th17 differentiation

Murine splenic T cells were isolated using Pan T Cell Isolation Kit II (Miltenyi Biotec, Bergisch Gladbach, Germany). Two hundred thousand T cells per well were cultured in 96-well plates in the presence of T Cell Activation/Expansion Kit (Miltenyi Biotec) for 2 weeks. The medium was supplemented twice per week with the following recombinant cytokines: mouse recombinant IL-6 (20 ng/mL), IL-1β (20 ng/mL), and IL-23 (40 ng/mL) for the IL-23 dependent Th17 cell condition; IL-6 (20 ng/mL) and TGFβ1 (3 ng/mL) for the IL-23 independent Th17 cell condition; or IL-23 (40 ng/mL) for the IL-23 only Th17 cell condition. The cytokines listed were purchased from BioLegend (Supplemental Table 5). Afterward, the cultured cells were processed for flow cytometry analysis.

### Western blotting

Murine epidermis was lysed using RIPA buffer (Wako) containing phosphatase-and protease-inhibitor cocktail (Nacalai tesque). Protein lysates were separated by 10% SuperSepTM Ace (Wako), transferred onto polyvinylidene difluoride membranes (0.45 μm, Merck, Darmstadt, Germany) by Trans-Blot Turbo Transfer System (Bio-Rad, Hercules, CA). The membrane was blocked with 5% bovine serum albumin and subjected to immunoblotting targeting the indicated proteins overnight 4℃, followed by the application of HRP-conjugated secondary antibody for one hour at room temperature (Supplemental Table 2). WB stripping solution (Nacalai tesque) was used to remove the antibodies for further evaluation.

### Inhibition of mTOR

Rapamycin (4 mg/kg, Sanxin Chempharma, Hebei, China), JR-AB2-011 (400 μg/kg, MedChemExpress, Monmouth Junction, NJ) and vehicle were applied intraperitoneally to mice once daily for 14 consecutive days. Rapamycin and JR-AB2- 011 were dissolved in 100% DMSO and diluted with 40% Polyethylene Glycol 300 (Wako), 5% Tween-80 (Sigma-Aldrich, St. Louis, MO) and 45% saline in sequence. For topical application, 0.2% rapamycin gel and vehicle gel were prepared by the pharmaceutical department as previously described (Wataya-Kaneda et al., 2017). Rapamycin gel was applied to the left ear, and vehicle gel to the right ear (10mg/ear) of Sema4AKO, once daily for 14 consecutive days. To analyze the preventive effectiveness of rapamycin in an IMQ-induced murine model of psoriatic dermatitis, Sema4AKO mice were administered either vehicle or rapamycin intraperitoneally from Day 0 to Day 17, and IMQ was topically applied to both ears for 4 days starting on Day 14. Then, on Day 18, ears were collected for further analysis.

### Data processing of single-cell RNA-sequencing and bulk RNA-sequencing

The raw count matrix data from the previously reported single-cell RNA- sequencing data from GSE220116 (Kim et al., 2023) were imported into Scanpy (1.9.6) using Python for further analyses. For each sample, cells and genes meeting the following criteria were excluded: cells expressing over 200 genes (sc.pp.filter_cells), cells with a high proportion of mitochondrial genes (> 5%), and genes expressed in fewer than 3 cells (sc.pp.filter_genes). Counts were normalized using sc.pp.normalize_per_cell, logarithmized (sc.pp.log1p), and scaled (sc.pp.scale). Highly variable genes were selected using sc.pp.filter_genes_dispersion with the options min_mean = 0.0125, max_mean = 2.5, and min_disp = 0.7.

Principal Component analysis was conducted with sc.pp.pca, selecting the 1st to 50th Principal Components for embedding and clustering. Neighbors were calculated with batch effect correction by BBKNN (Polański et al., 2020). Embedding was performed by sc.tl.umap, and cells were clustered using sc.tl.leiden. Some cells were further subclassified in a similar manner. The data was integrated into an h5ad file, which can be visualized in Cellxgene VIP (K. Li et al., 2022). We then performed differential analysis between two groups of cells to identify differential expressed genes using Welch’s t-test. Multiple comparisons were controlled using the Benjamini- Hochberg procedure, with the false discovery rate set at 0.05 and significance defined as *padj* < 0.05.

Bulk RNA-sequencing data from GSE121212 (Tsoi et al., 2019) were re- analyzed using RaNAseq (Prieto & Barrios, 2019) for differential expression in Ctl versus psoriatic NL. Our analysis included normalization, differential gene expression (defining significance as *padj* < 0.05), and focused on Gene Ontology biological process analysis. The gene expression was calculated with the transcripts per million values.

### Statistical analysis

GraphPad Prism 10 software (GraphPad Software, La Jolla, CA) was used for all statistical analyses except for RNA-sequencing. Mann-Whitney test was used for two-group comparisons, while Kruskal-Wallis test followed by Dunn’s multiple comparisons test was applied for comparison among three or more groups. Statistical significance was defined as *p* < 0.05 (*), *p* < 0.01 (**), and *p* < 0.001 (***). In the analysis of RNA-sequencing data, statistical significance was defined as *padj* < 0.01 (**), and *padj* < 0.001 (***).

### Study approval

All experiments involving human specimens were in accordance with the Declaration of Helsinki and were approved by the Institutional Review Board in Osaka University Hospital (20158-6). Written informed consent was obtained from all participants. All murine experiments were approved by Osaka University Animal Experiment Committee (J007591-013) and all procedures were conducted in compliance with the Guidelines for Animal Experimentation established by Japanese Association for Laboratory Animal Science.

### Data availability

The single-cell RNA-sequencing datasets generated by Kim et al. (Kim et al., 2023) and Tsoi et al. (Tsoi et al., 2019) used in this study are available in the NCBI Gene Expression Omnibus under accession codes GSE220116 and GSE121212, respectively. Values for all data points in graphs are reported in the Source data.

## Supporting information

Key Resources Table

## Acknowledgments

We express our gratitude to all patients for their participation in this study.

## SUPPLEMENTARY MARERIALS

**Figure 1-figure supplement 1:**
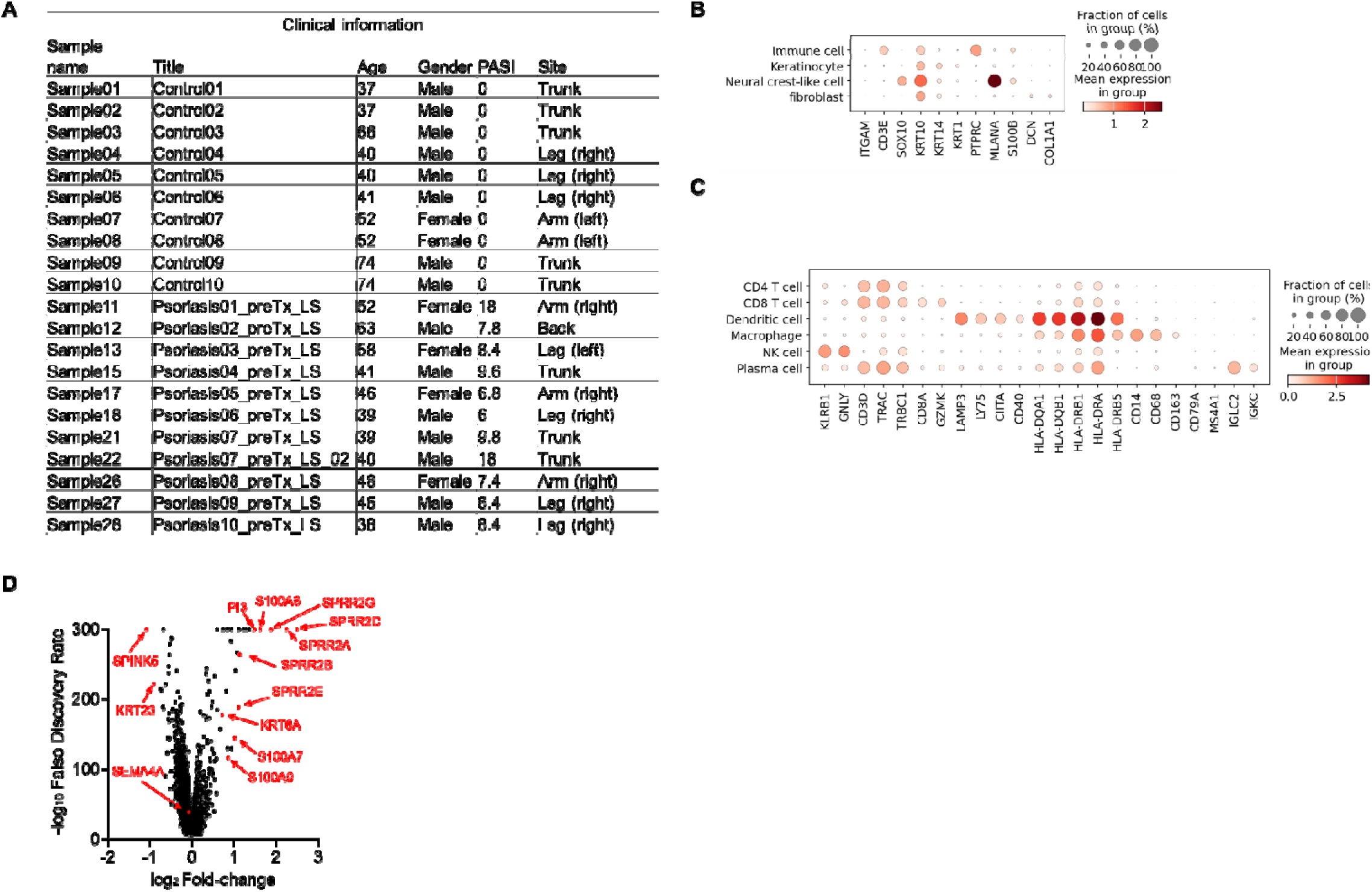
Sema4A is downregulated in the keratinocytes of lesional psoriasis in the single-cell RNA-sequencing data. (A) Sample information for specimens from Ctl and psoriatic L (GSE220116). (B and C) Clusters of cells were identified by their expression patterns of signature genes. (D) The volcano plot displays changes in gene expression in psoriatic L compared to Ctl. Figure 1-figure supplement 1-source data 1 Excel file containing quantitative data for Figure 1-figure supplement 1.

**Figure 1-figure supplement 2:**
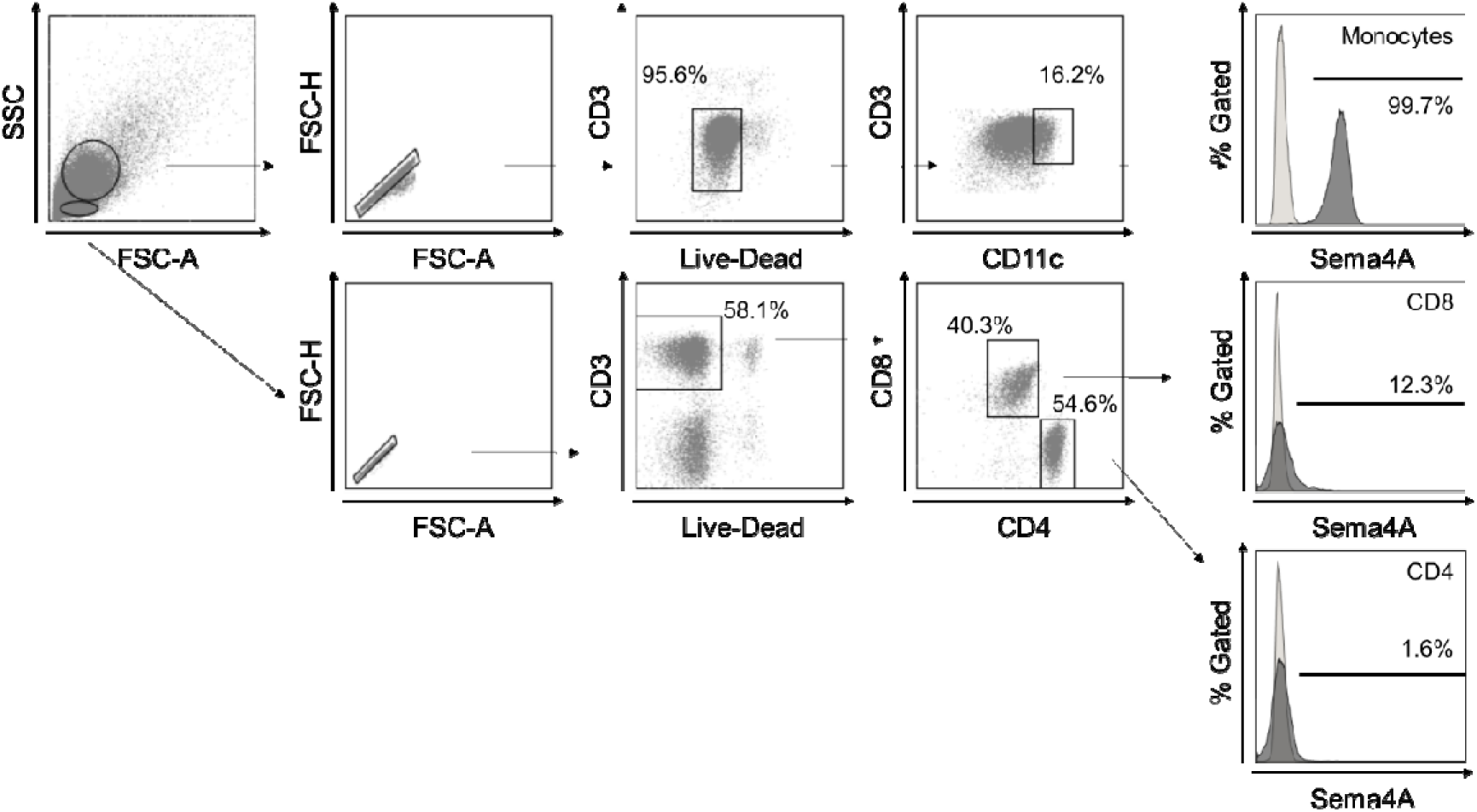
Gating strategy in flow cytometry. Gating strategy for human Sema4A expression in blood cells. Large and small cells were distinguished using forward scatter (FCS) and side scatter (SSC) in a dot plot panel, with dead cells being excluded. Monocytes were defined within the live large cell population as CD11c positive. CD4 and CD8 T cells were identified within the live small cell population as CD3 positive CD4 positive and CD3 positive CD8 positive populations, respectively. The empty histogram represents the flow cytometry minus one control for Sema4A.

**Figure 2-figure supplement 1:**
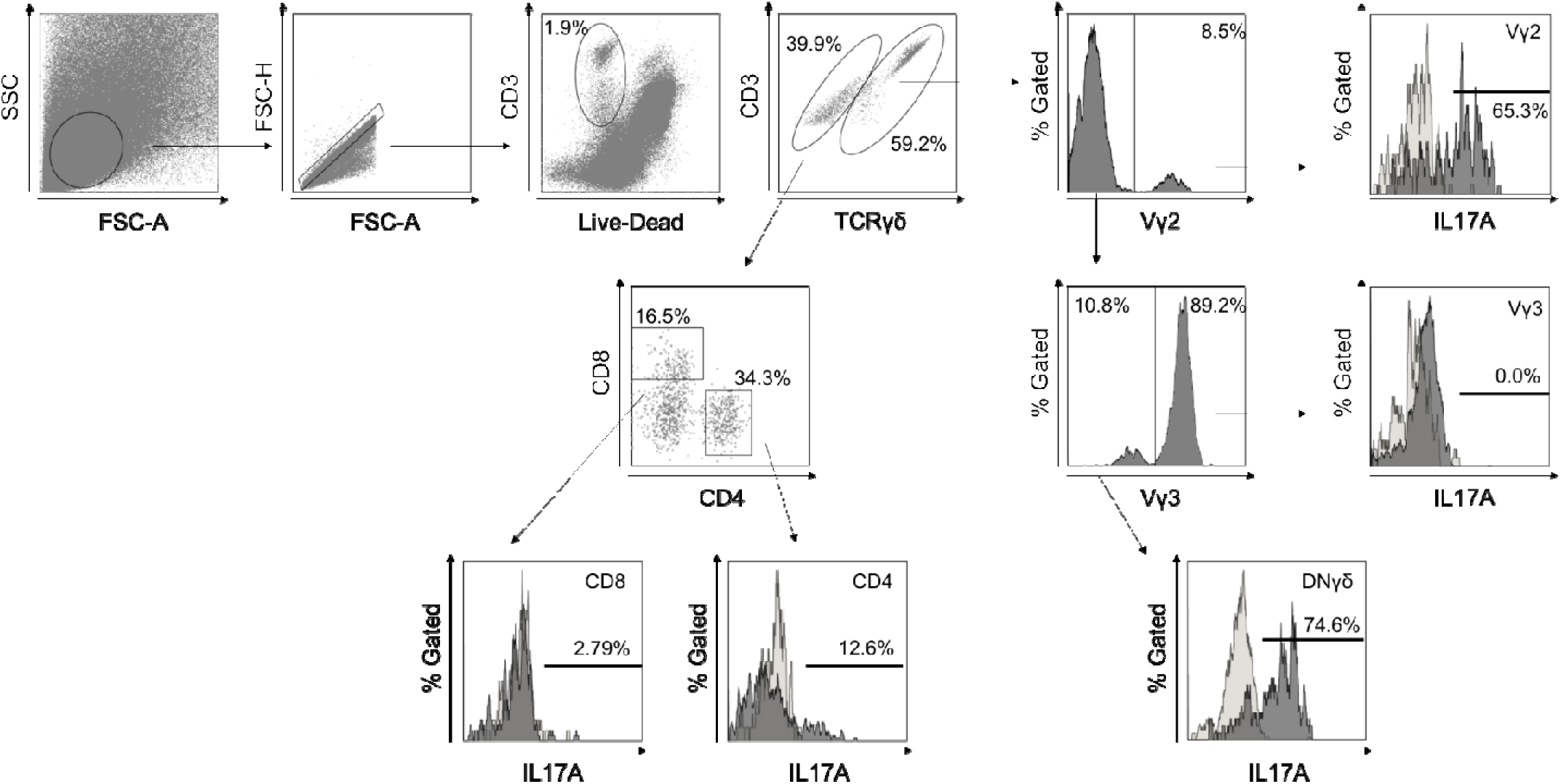
Gating strategy in flow cytometry. Gating strategy for murine T cells infiltrating the epidermis and dermis. After excluding dead cells, TCRγδ positive T cells were evaluated for the expression of Vγ2. TCRγδ positive Vγ2 negative population was further assessed the expression of Vγ3. The CD3 positive TCRγδ negative population was evaluated for the expression of CD4 and CD8. Each population was analyzed for cytokine production. The empty histogram represents the isotype control for IL-17A.

**Figure 2-figure supplement 2:**
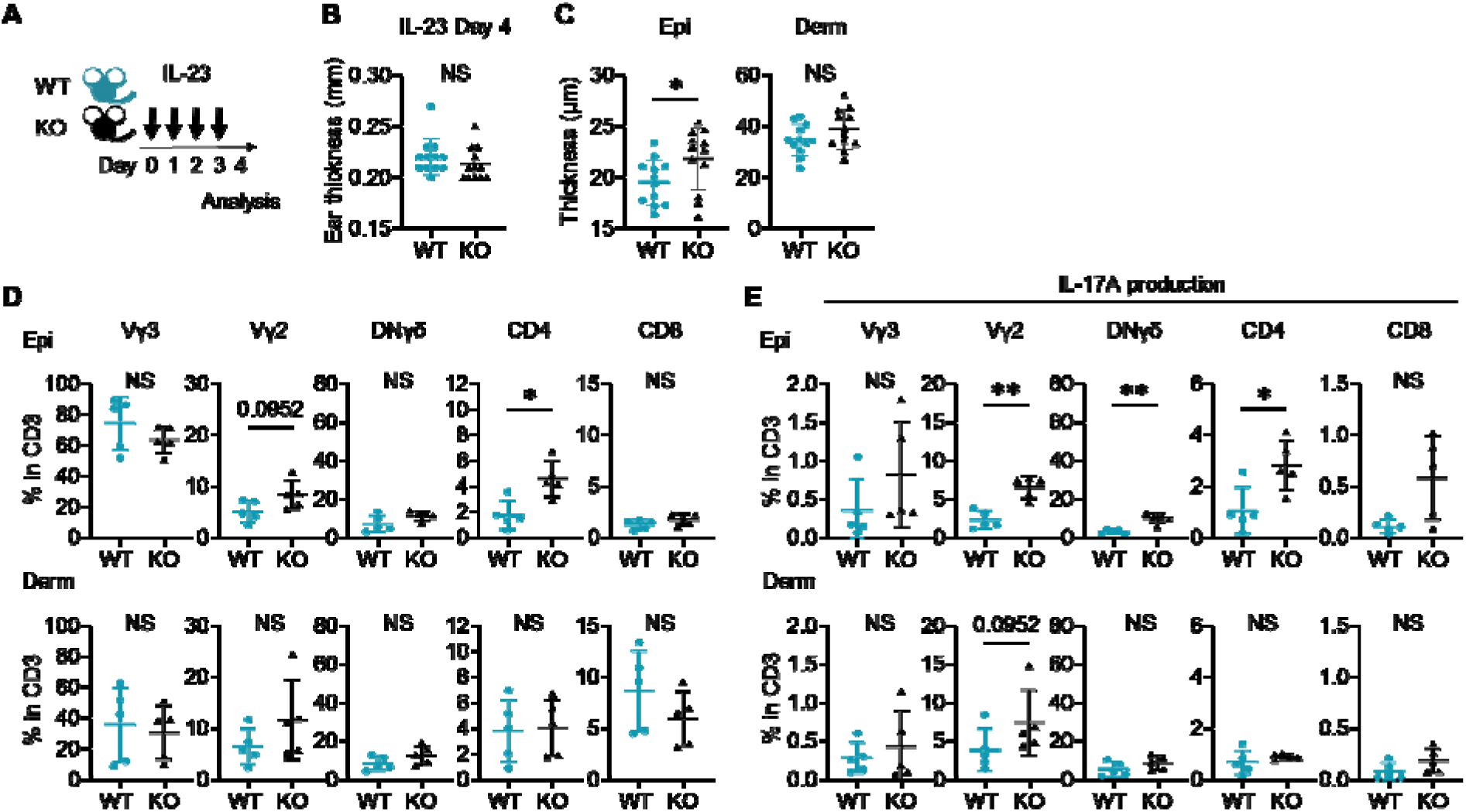
IL-23-mediated psoriasis-like dermatitis is augmented in Sema4AKO mice. (A) An experimental scheme involved intradermally injecting 20 μl of phosphate- buffered saline containing 500 ng of recombinant mouse IL-23 into both ears of WT mice and KO mice for 4 consecutive days. Samples for following analysis were collected on Day 4. (B and C) Ear thickness (B) and Epi and Derm thickness (C) of WT mice and KO mice on Day 4 (*n* = 12 per group). (D and E) The percentages of Vγ3, Vγ2, DNγδ, CD4, and CD8 T cells (D) and those with IL-17A production (E) in CD3 fraction in the Epi (top) and Derm (bottom) of WT and KO ears (*n* = 5 per group). Each dot represents the average of 4 ear specimens. B-E: \**p* < 0.05, \*\**p* < 0.01. NS, not significant. Figure 2-figure supplement 2-source data 1 Excel file containing quantitative data for Figure 2-figure supplement 2.

**Figure 3-figure supplement 1:**
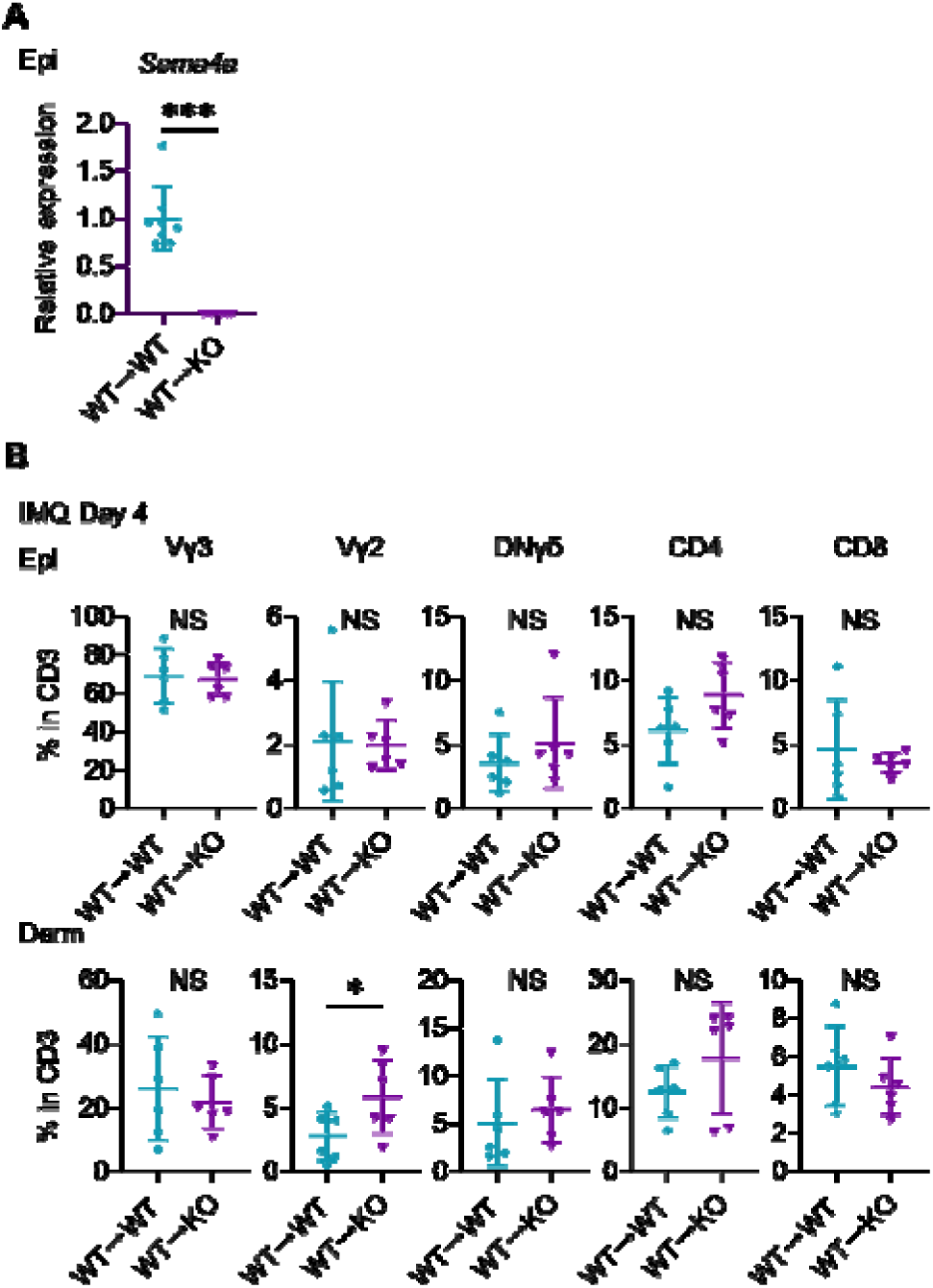
T cells’ fractions infiltrating in the chimeric mice ear. (A) *Sema4a* expression in the Epi of WT→ WT mice and WT→ KO mice (*n* = 8for WT→ WT, *n* = 7 for WT→ KO). (B) The percentages of Vγ3, Vγ2, DNγδ, CD4, and CD8 T cells in CD3 fraction from IMQ Day 4 Epi (top) and Derm (bottom) of the ears from WT→ WT mice and WT→ KO mice (*n* = 6 per group). Each dot represents the average of 4 ear specimens. A-B: \**p* < 0.05, \*\*\**p* < 0.001. NS, not significant. Figure 3-figure supplement 1-source data 1 Excel file containing quantitative data for Figure 3-figure supplement 1.

**Figure 4-figure supplement 1:**
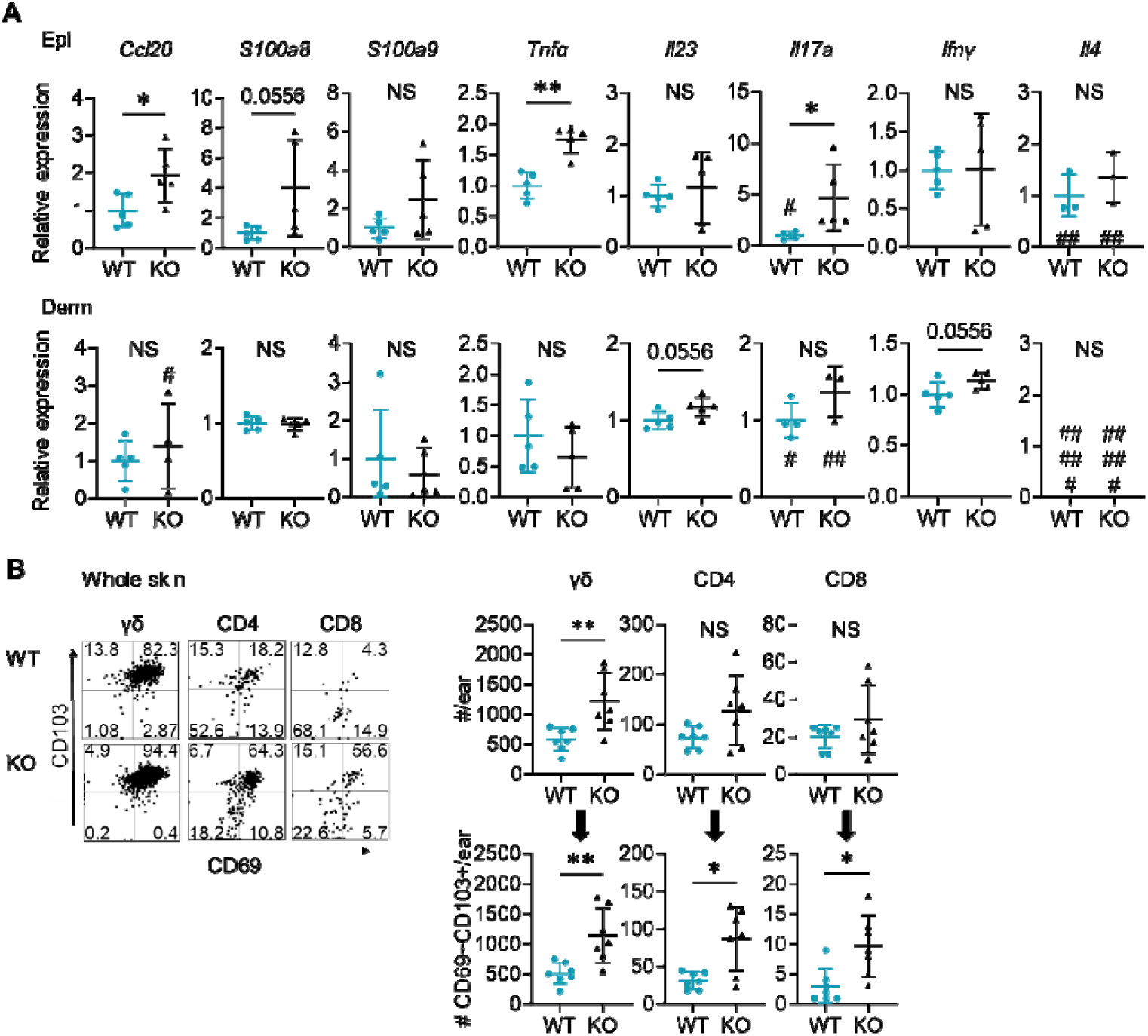
Naive Sema4AKO skin shows upregulation of psoriasis related genes and an increase in resident memory T cells. (A) Relative expression of psoriasis-associated genes in Epi (top) and Derm (bottom) of WT mice and KO mice (*n* = 5 per group, #: not detected). (B) Representative dot plots showing CD69 and CD103 expression in the indicated T cell fractions from whole skin. The graphs show T cell counts per ear (top) and those with resident memory phenotype (bottom) (*n* = 7 per group). Each dot represents the average of 4 ear specimens. A-B: \**p* < 0.05, \*\**p* < 0.01. NS, not significant. Figure 4-figure supplement 1-source data 1 Excel file containing quantitative data for Figure 1-figure supplement 1.

**Figure 4-figure supplement 2:**
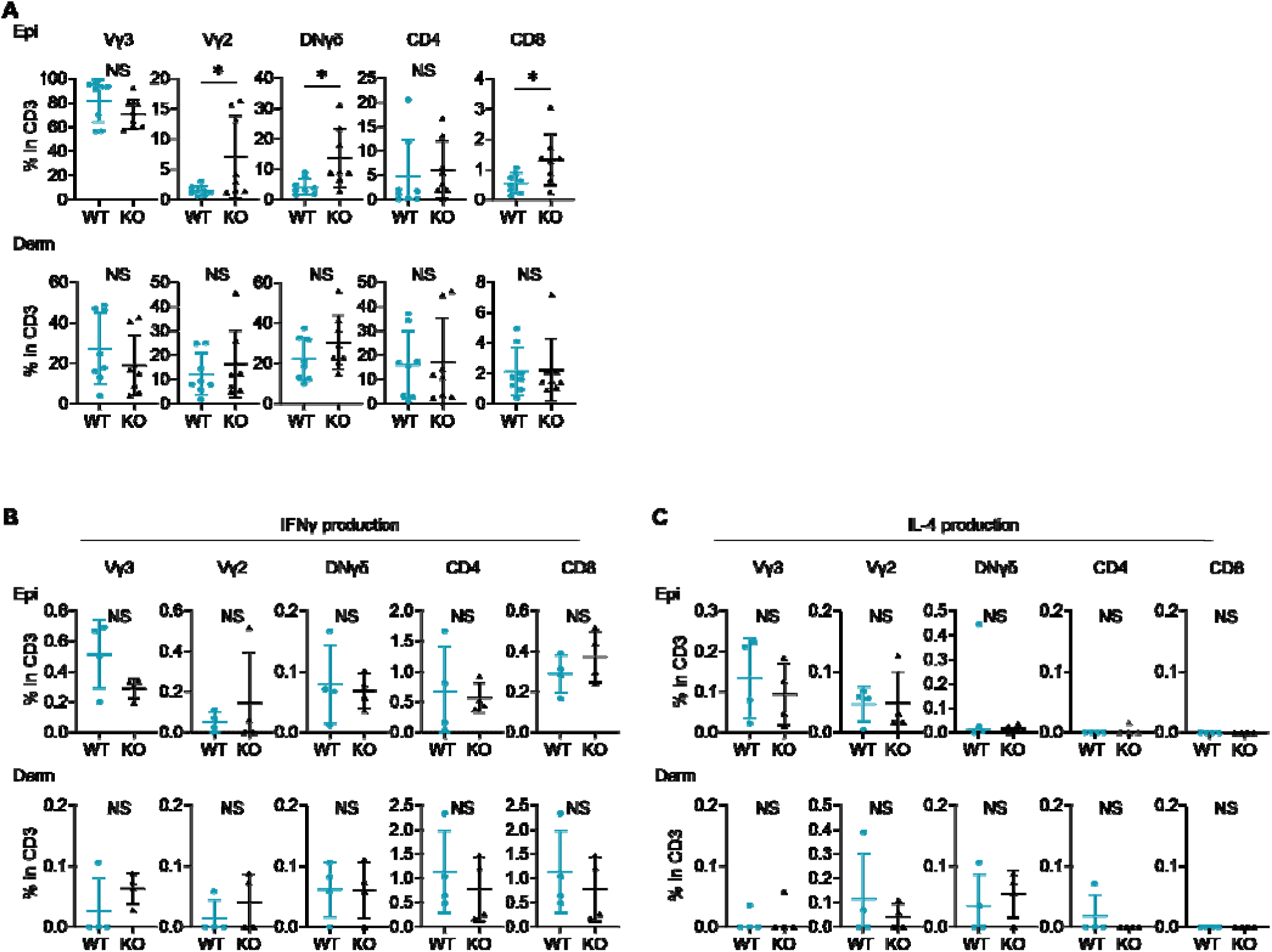
Expression of IFNγ and IL-4is comparable between naive WT and Sema4AKO skin. (A) The percentages of Vγ3, Vγ2, DNγδ, CD4, and CD8 T cells in CD3 fraction from naive WT and KO mice. (B and C) The graphs presenting the percentages of IFNγ (B) and IL-4 (C) -producing Vγ2, DNγδ, CD4, and CD8 T cells in CD3 fraction in the Epi (top) and Derm (bottom) of naive WT mice and KO mice (*n* = 4 per group). A-C: Each dot represents the average of 4 ear specimens. \**p* < 0.05. NS, not significant. Figure 4-figure supplement 2-source data 1 Excel file containing quantitative data for Figure 4-figure supplement 2.

**Figure 4-figure supplement 3:**
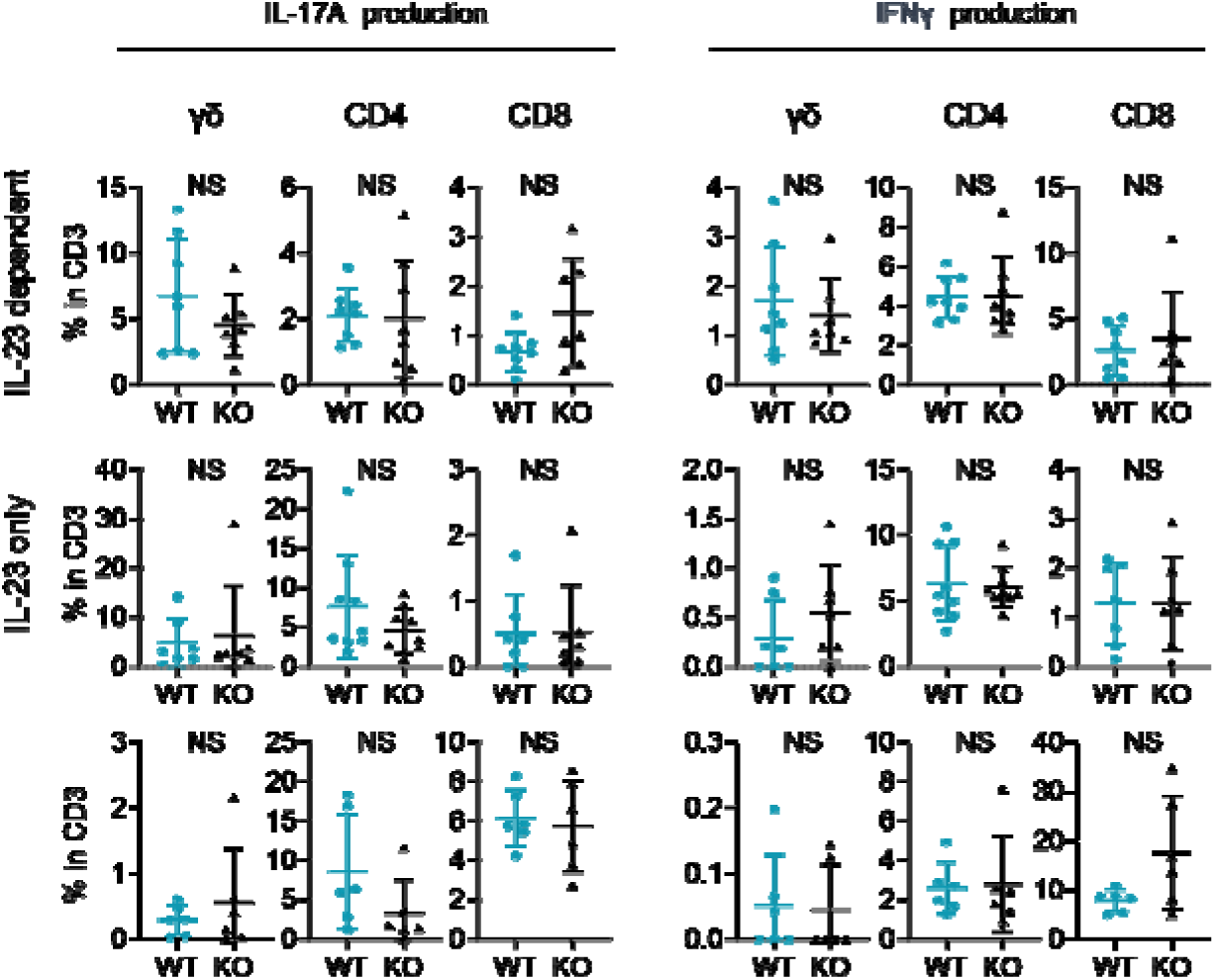
Comparable T17differentiation potential under Th17-skewing conditions between WT mice and Sema4AKO mice. Splenic T cells were cultured for 2 weeks, followed by flow cytometry analysis. The accumulated data display the percentages of IL-17A-producing (right) and IFNγ- producing (left) γδ, CD4, and CD8 T cells within CD3 fraction under various conditions: IL-23 dependent Th17-skewing condition (top), IL-23 only Th17-skewing condition (middle), and IL-23 independent Th17-skewing condition (bottom) . NS, not significant. Figure 4-figure supplement 3-source data 1 Excel file containing quantitative data for Figure 4-figure supplement 3.

**Figure 5-figure supplement 1:**
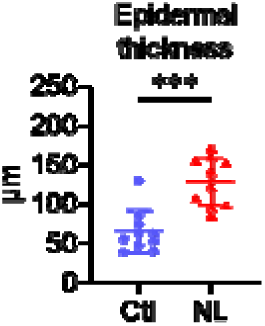
The epidermis of psoriatic non-lesion is thicker than that of control skin. Epidermal thickness of Ctl and psoriatic NL (*n* = 10 per group). \*\*\**p* < 0.001. Figure 5-figure supplement1-source data1 Excel file containing quantitative data for Figure 5-figure supplement1.

**Figure 5-figure supplement 2:**
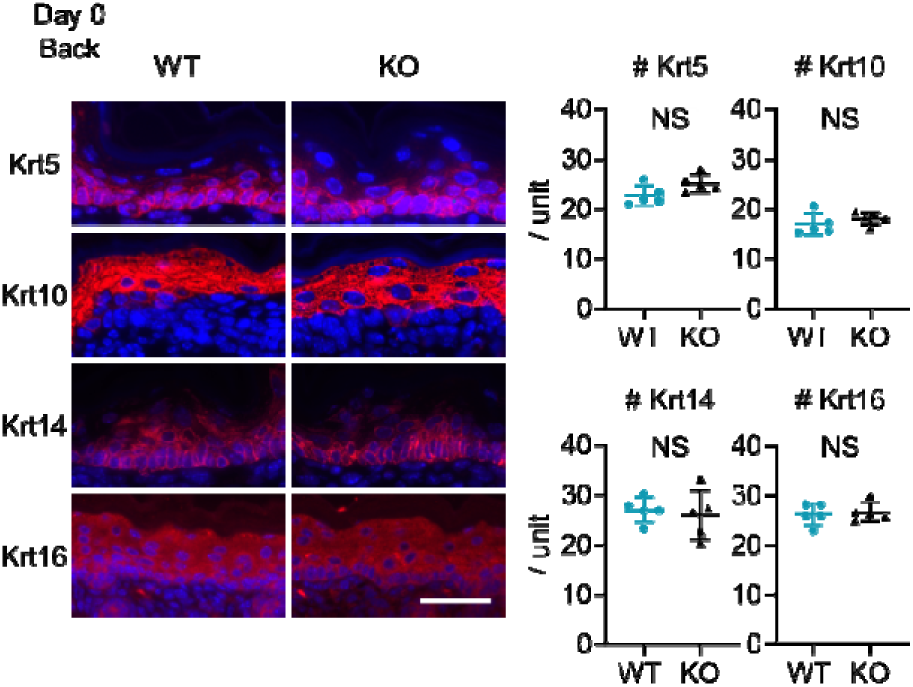
Upregulation of cytokeratin expression related to psoriasis is not detected at birth in Sema4AKO mice. Representative immunofluorescence pictures of Krt5, Krt10, Krt14, and Krt16 (red) overlapped with DAPI, and the accumulated graphs showing the numbers of Krt5, Krt10, Krt14, and Krt16 positive cells per 100 μm width (*n* = 5 per group) in the epidermis of Day 0 back. Scale bar = 50 μm. Each dot represents the average from 5 unit areas per sample. NS, not significant. Figure 5-figure supplement 2-source data 1 Excel file containing quantitative data for Figure 5-figure supplement 2.

**Figure 7-figure supplement 1:**
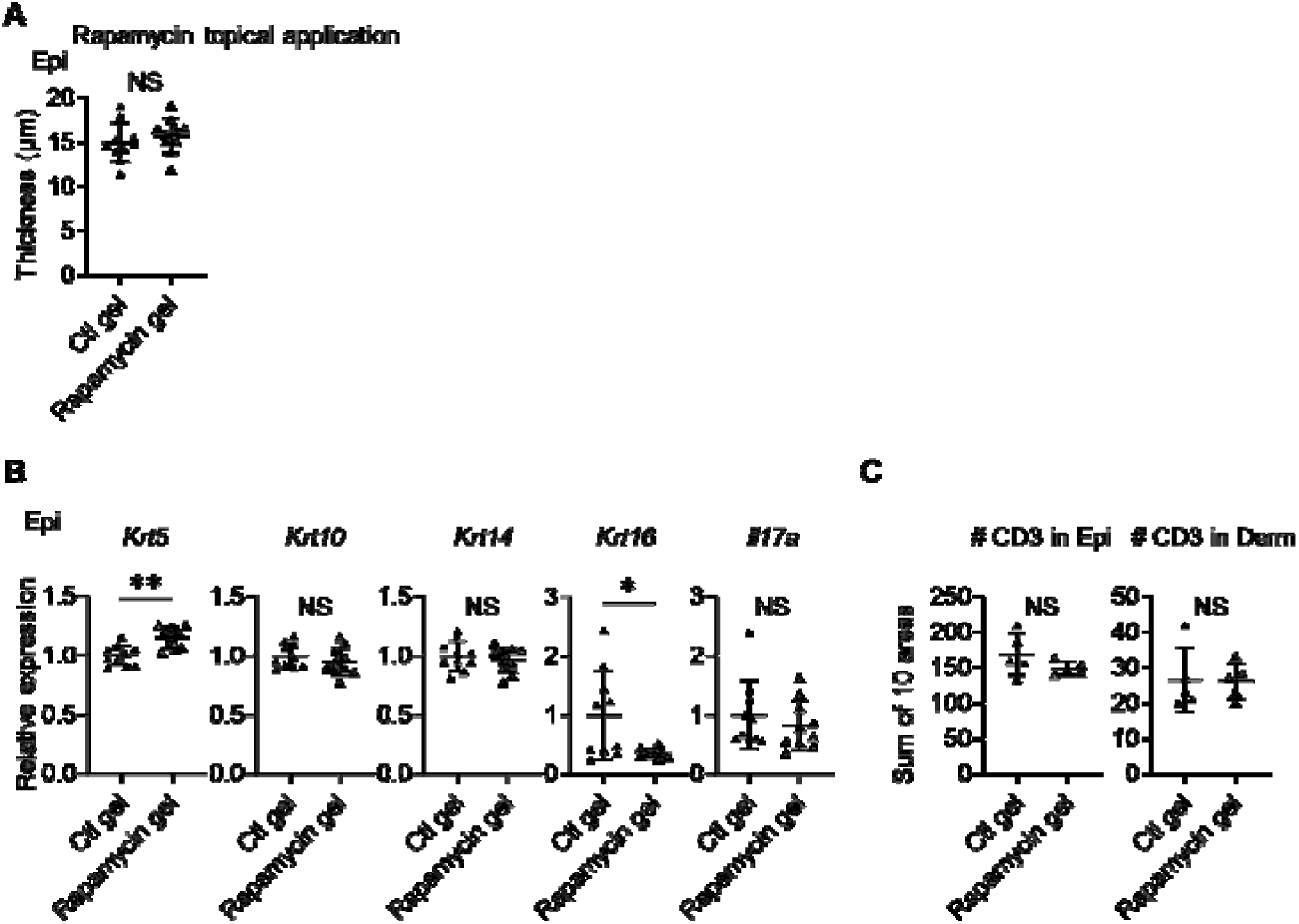
Topical application of rapamycin gel yields partially similar results to intraperitoneal treatment. (A) Comparison of Epi thickness between vehicle (Ctl) gel-treated right ears and rapamycin gel-treated left ears of Sema4AKO mice (*n* = 10 per group). (B) Relative expression of keratinocyte differentiation markers and *Il17a* in Sema4AKO Epi under Ctl gel or rapamycin gel treatments (*n* = 5 per group). (C) The number of T cells in the Epi (left) and Derm (right), under Ctl gel or Rapamycin gel treatments (*n* = 5 per group). Each dot represents the sum of numbers from 10 unit areas across 3 specimens. A-C: \**p* < 0.05, \*\**p* < 0.01. NS, not significant. Figure 7-figure supplement 1-source data 1 Excel file containing quantitative data for Figure 7-figure supplement 1.

**Figure 7-figure supplement 2:**
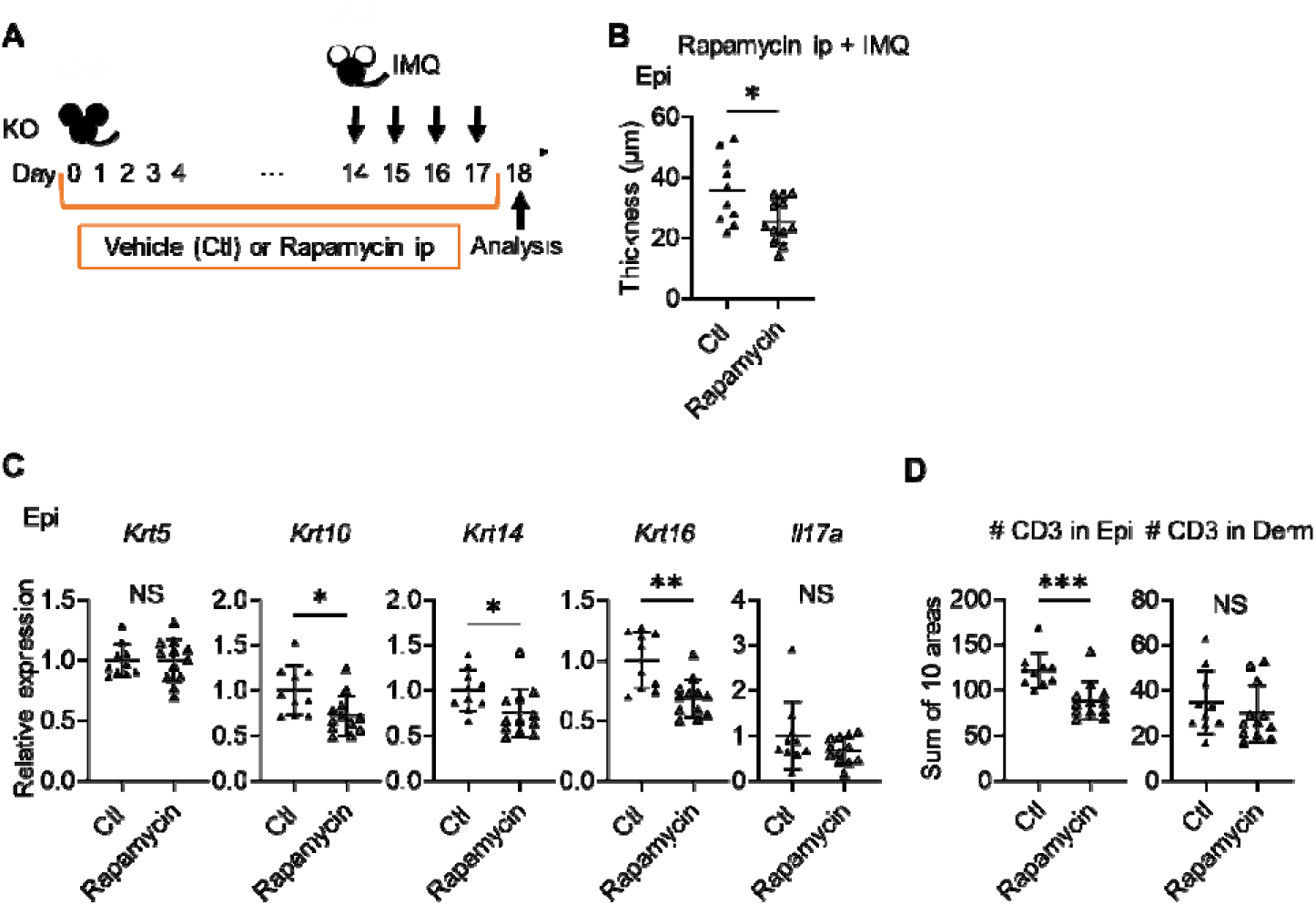
Rapamycin treatment reduced the epidermal swelling observed in IMQ-treated Sema4AKO mice. (A) Experimental scheme. (B) The Epi thickness on Day 18. (*n* = 10 for Ctl, *n* = 12 for Rapamycin). (C) Relative expression of keratinocyte differentiation markers and *Il17a* in Sema4AKO Epi (*n* = 10 for Ctl, *n* = 12 for Rapamycin). (D) The number of T cells in the Epi (left) and Derm (right), under Ctl or rapamycin and IMQ treatments (*n* = 10 for Ctl, *n* = 12 for Rapamycin). Each dot represents the sum of numbers from 10 unit areas across 3 specimens. A-C: \**p* < 0.05, \*\**p* < 0.01. NS, not significant. Figure 7-figure supplement 2-source data 1 Excel file containing quantitative data for Figure 7-figure supplement 2.

**Table S1.**
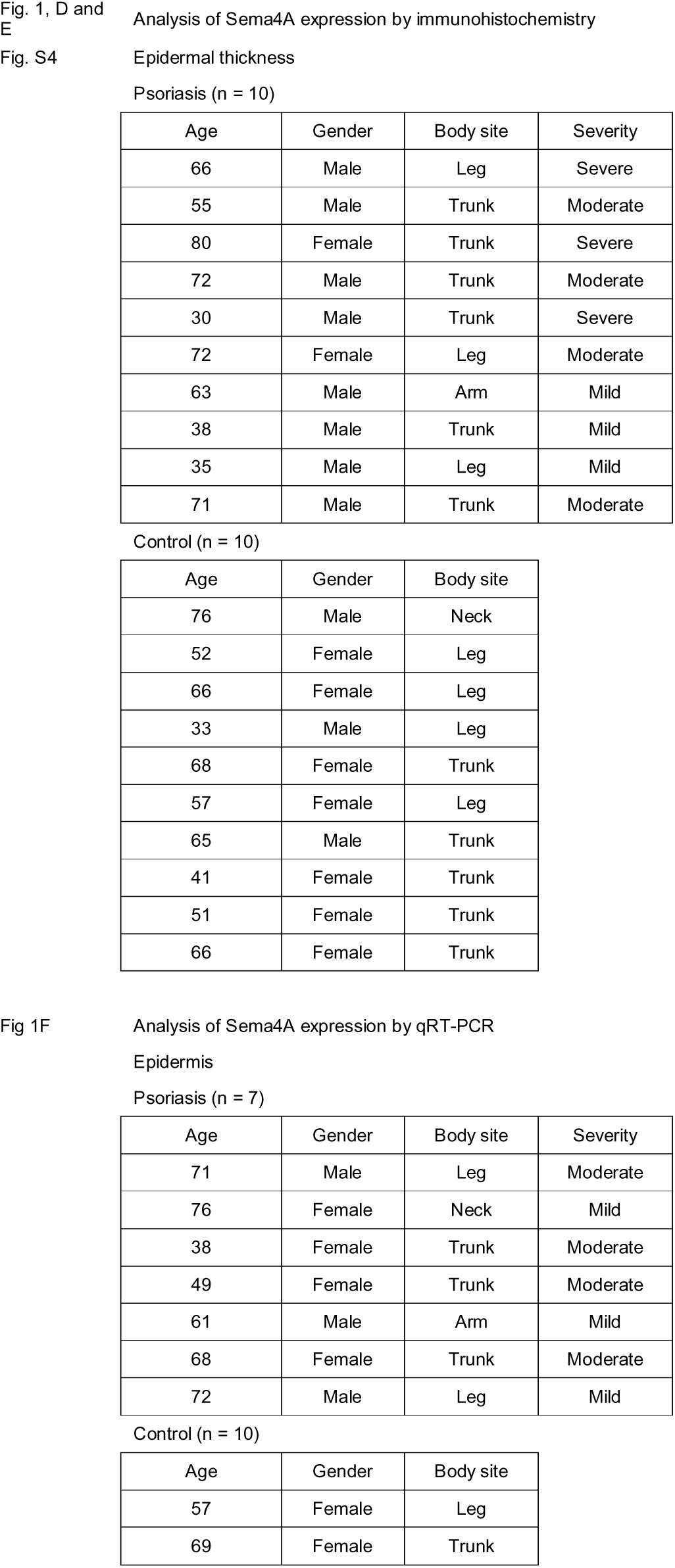
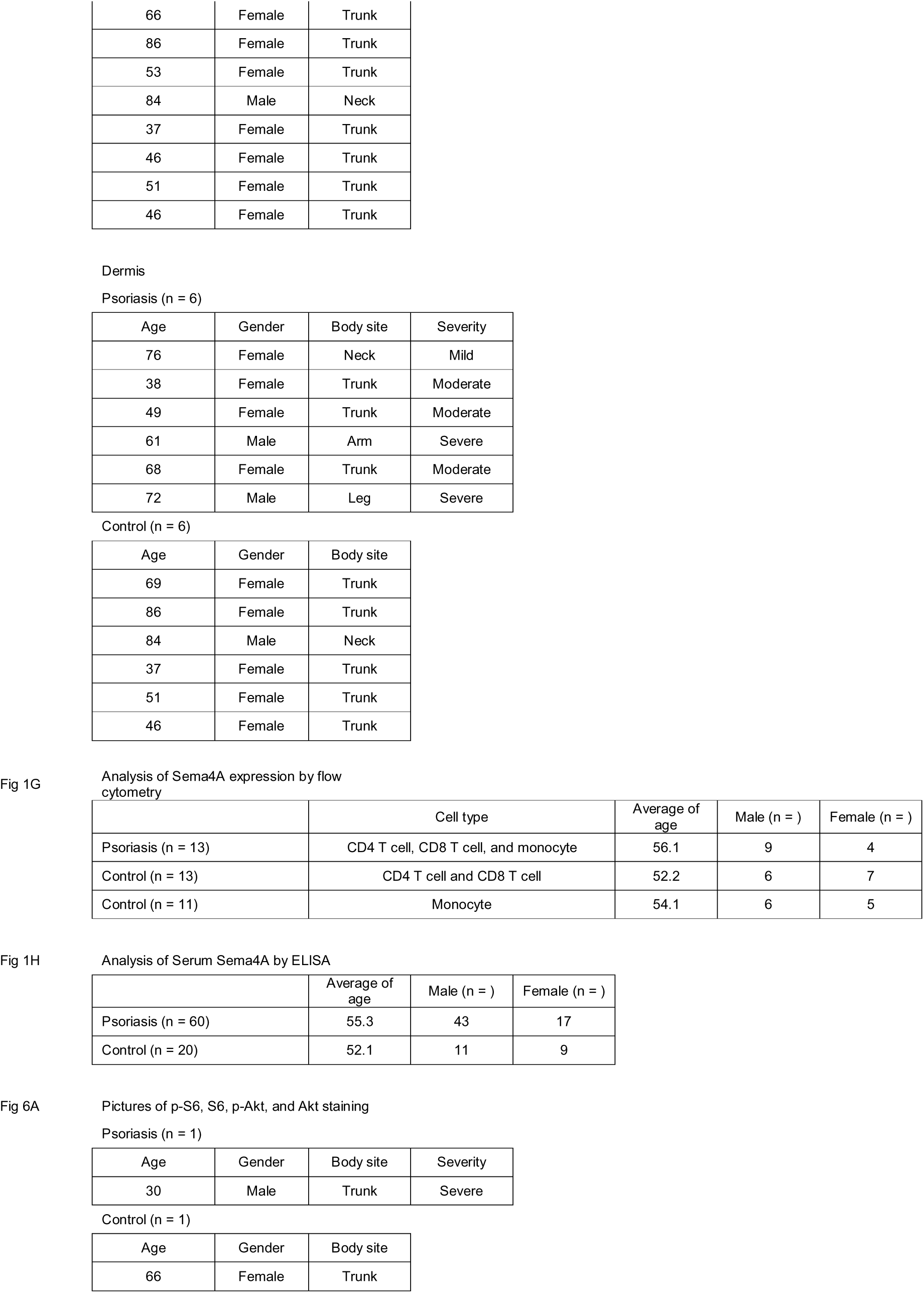
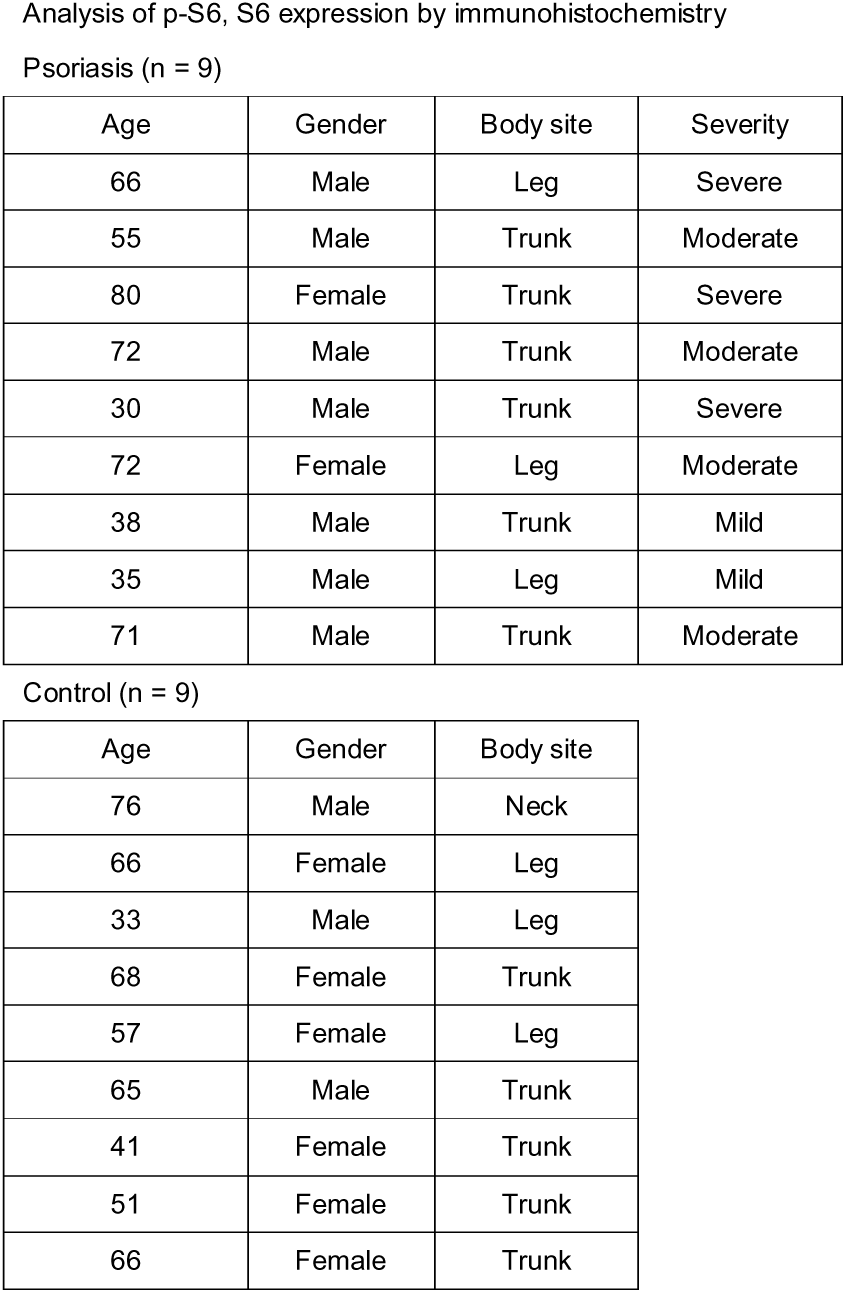
Patient information.

**Table S2.**
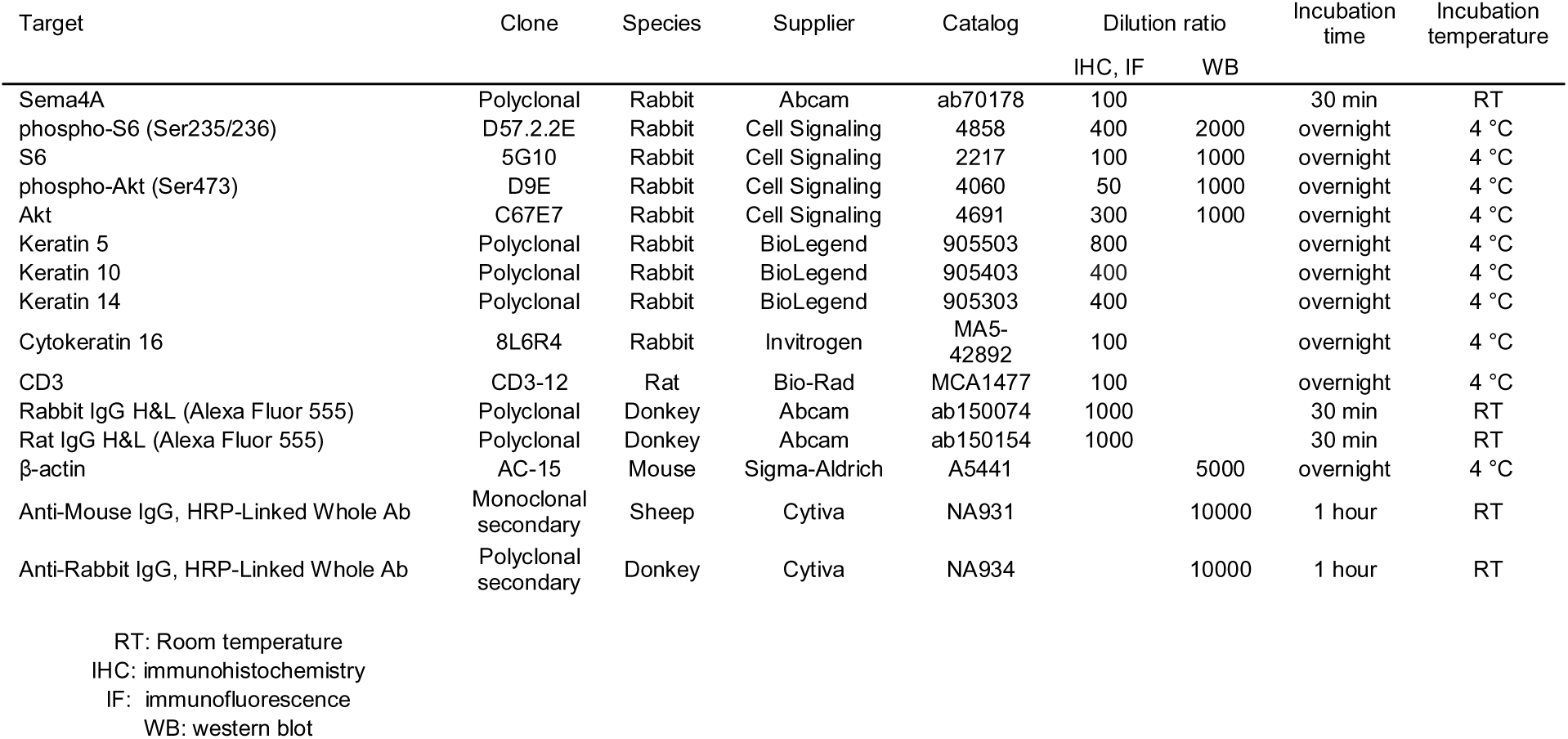
Antibodies used for immunohistochemical, immunofluorescence, and western blot analyses.

**Table S3.**
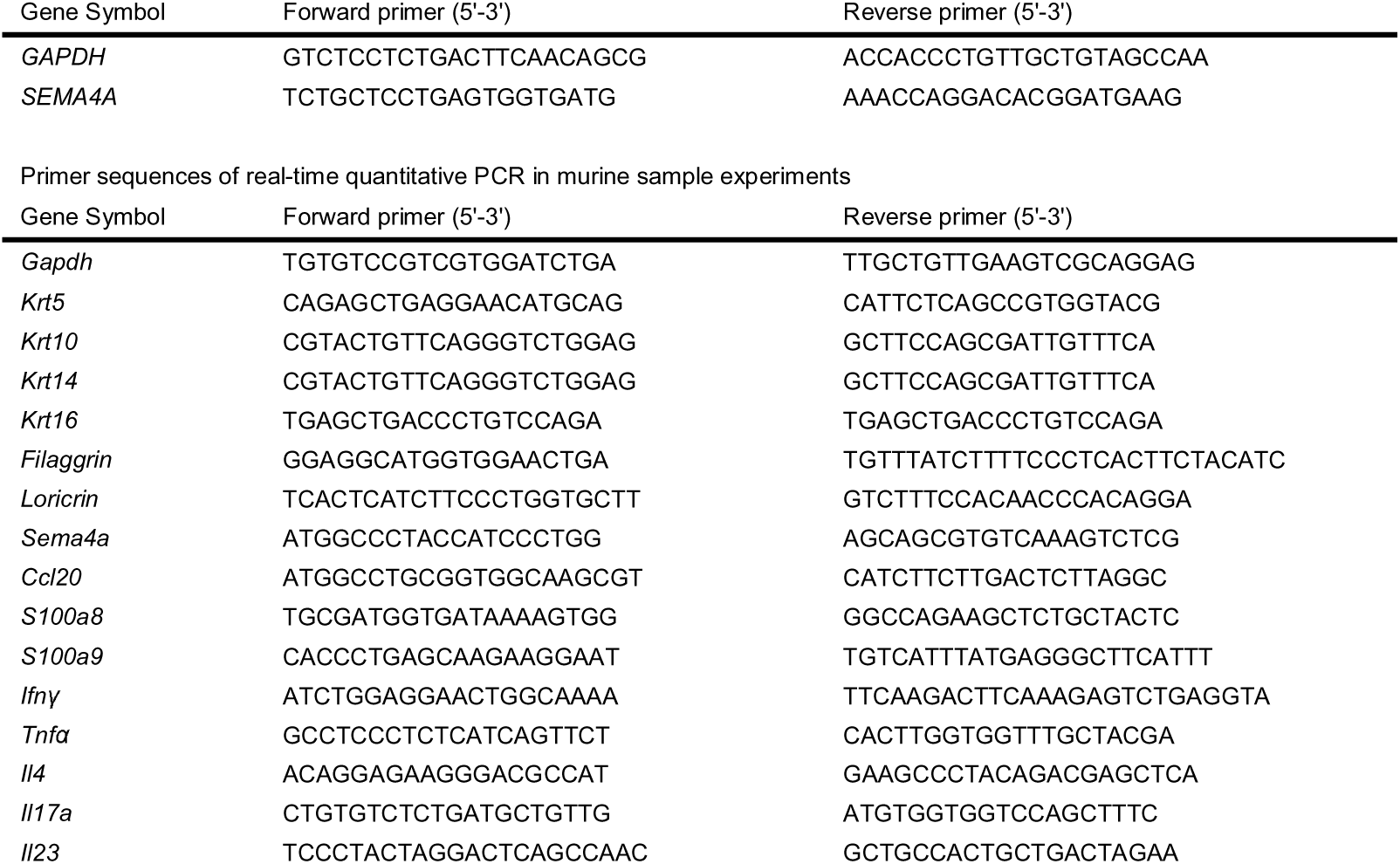
Primer sequences of real-time quantitative PCR used in human sample experiments.

**Table S4.** Antibodies used for flow cytometry analysis.

| Target | Clone | Species | Supplier | Catalog |
| --- | --- | --- | --- | --- |
| CD3 | OKT3 | Mouse | BioLegend | 317335 |
| CD4 | RPA-T4 | Mouse | BioLegend | 300512 |
| CD8a | RPA-T8 | Mouse | eBioscience | 47-0088-42 |
| CD11c | 3.9 | Mouse | BioLegend | 301628 |
| SEMA4A | 5E3/SEMA4A | Mouse | BioLegend | 148404 |
| CD3e | 145-2C11 | Armenian Hamster | BioLegend | 100328 |
| CD4 | GK1.5 | Rat | BioLegend | 100406 |
| CD8a | 53-6.7 | Rat | BioLegend | 100714 |
| CD16/32 | 93 | Rat | BioLegend | 101301 |
| CD69 | H1.2F3 | Armenian Hamster | BioLegend | 104514 |
| CD103 | 2E7 | Armenian Hamster | BioLegend | 121422 |
| TCR Vγ2 | UC3-10A6 | Armenian Hamster | BioLegend | 137705 |
| TCR Vγ3 | 536 | Syrian Hamster | BD Biosciences | 743241 |
| TCRγ $\delta$ | GL3 | Armenian Hamster | BioLegend | 118124 |
| IFNγ | XMG1.2 | Rat | BioLegend | 505813 |
| IL-4 | 11B11 | Rat | BD Biosciences | 562915 |
| IL-17A | TC11-18H10.1 | Rat | BioLegend | 506925 |

**Table S5.** Mouse recombinant cytokines.

| Cytokines | Supplier | Catalog |
| --- | --- | --- |
| IL-1β | BioLegend | 575102 |
| IL-6 | BioLegend | 575702 |
| IL-23 | BioLegend | 589002 |
| TGFβ1 | BioLegend | 763102 |

