## Supplementary figures and images for "Downregulation of Semaphorin 4A in keratinocytes reflects the features of non-lesional psoriasis"

### Figure 1-figure supplement 1

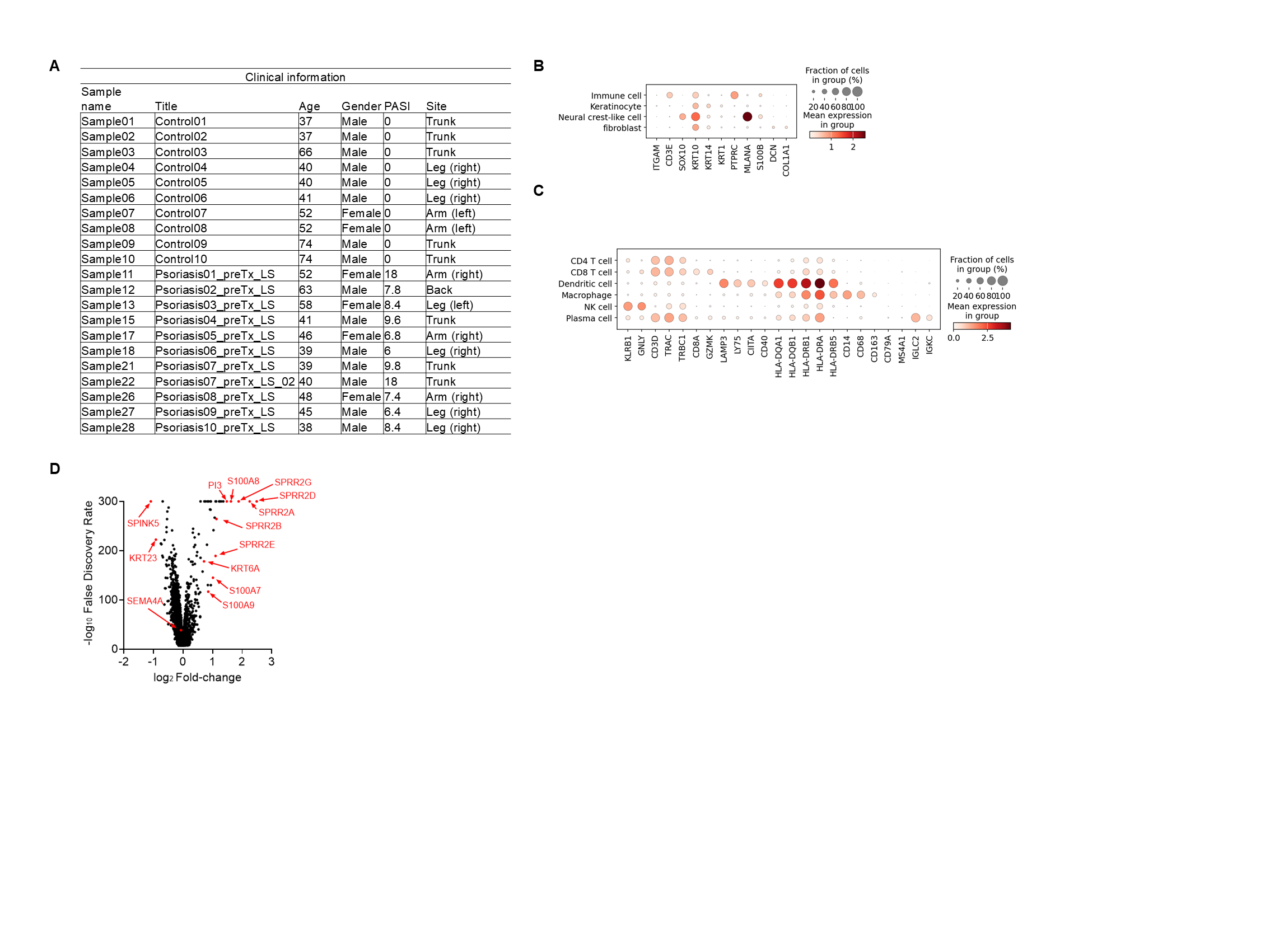

### Figure 1-figure supplement 2

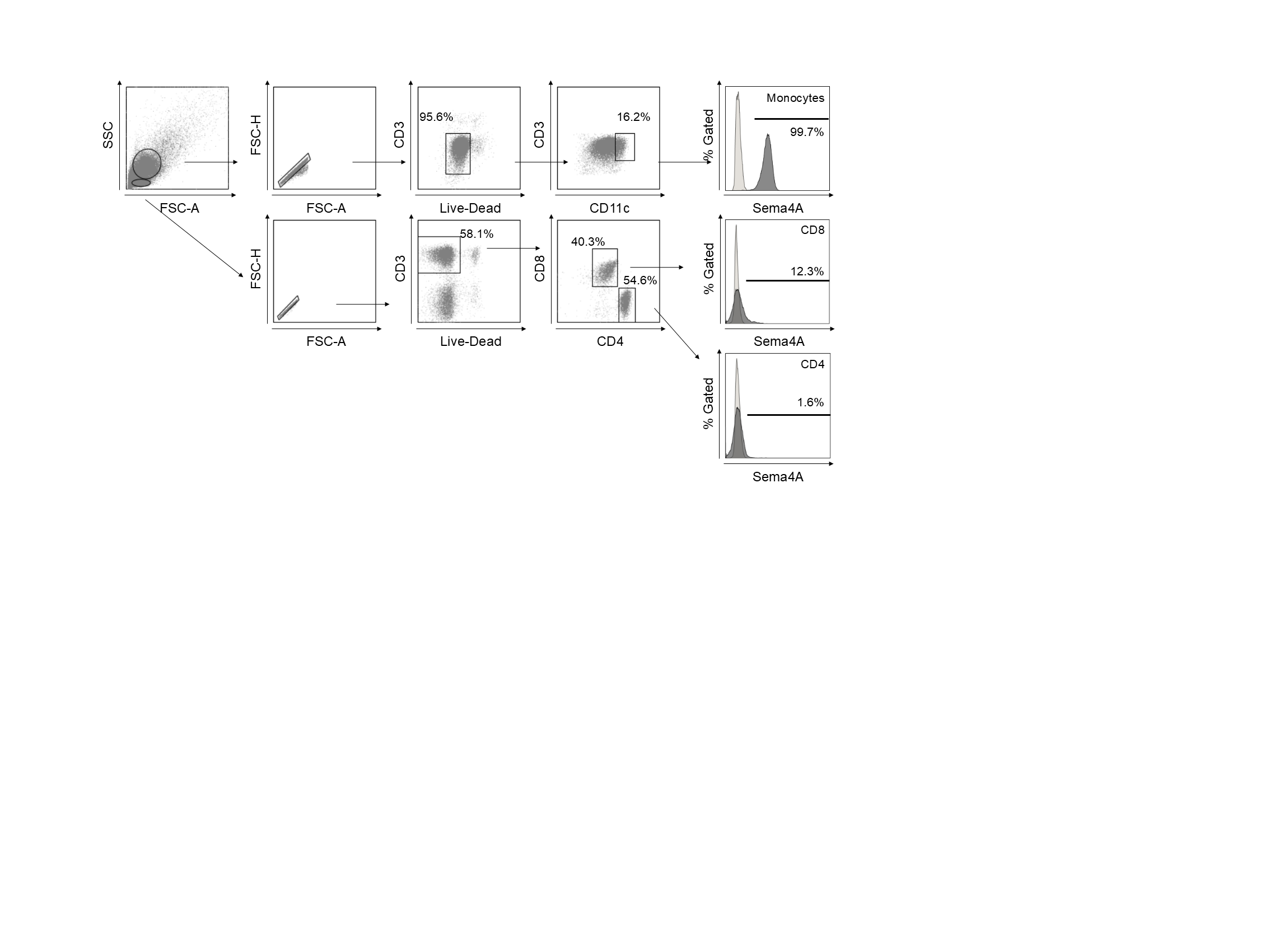

### Figure 2-figure supplement 1

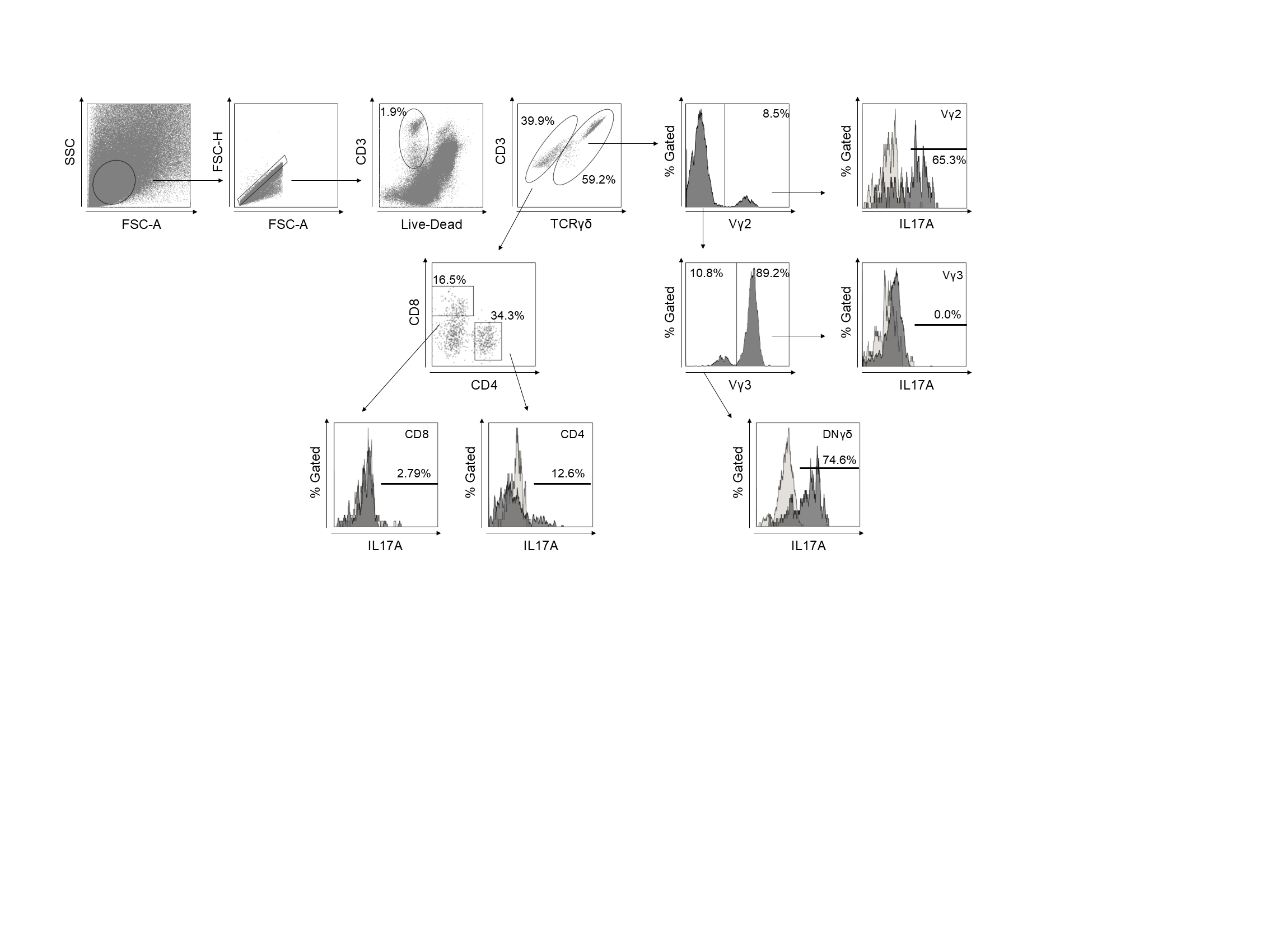

### Figure 2-figure supplement 2

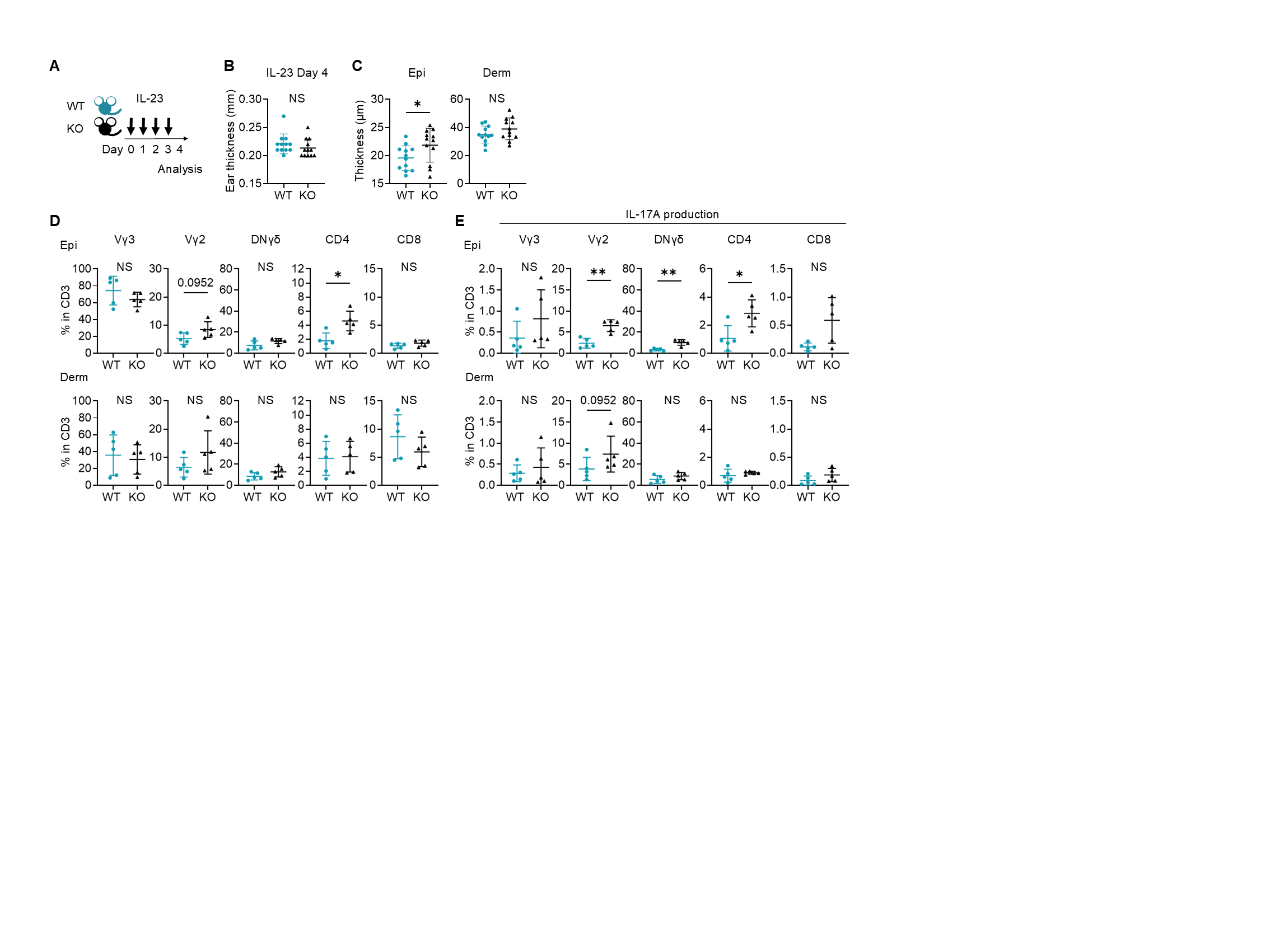

### Figure 3-figure supplement 1

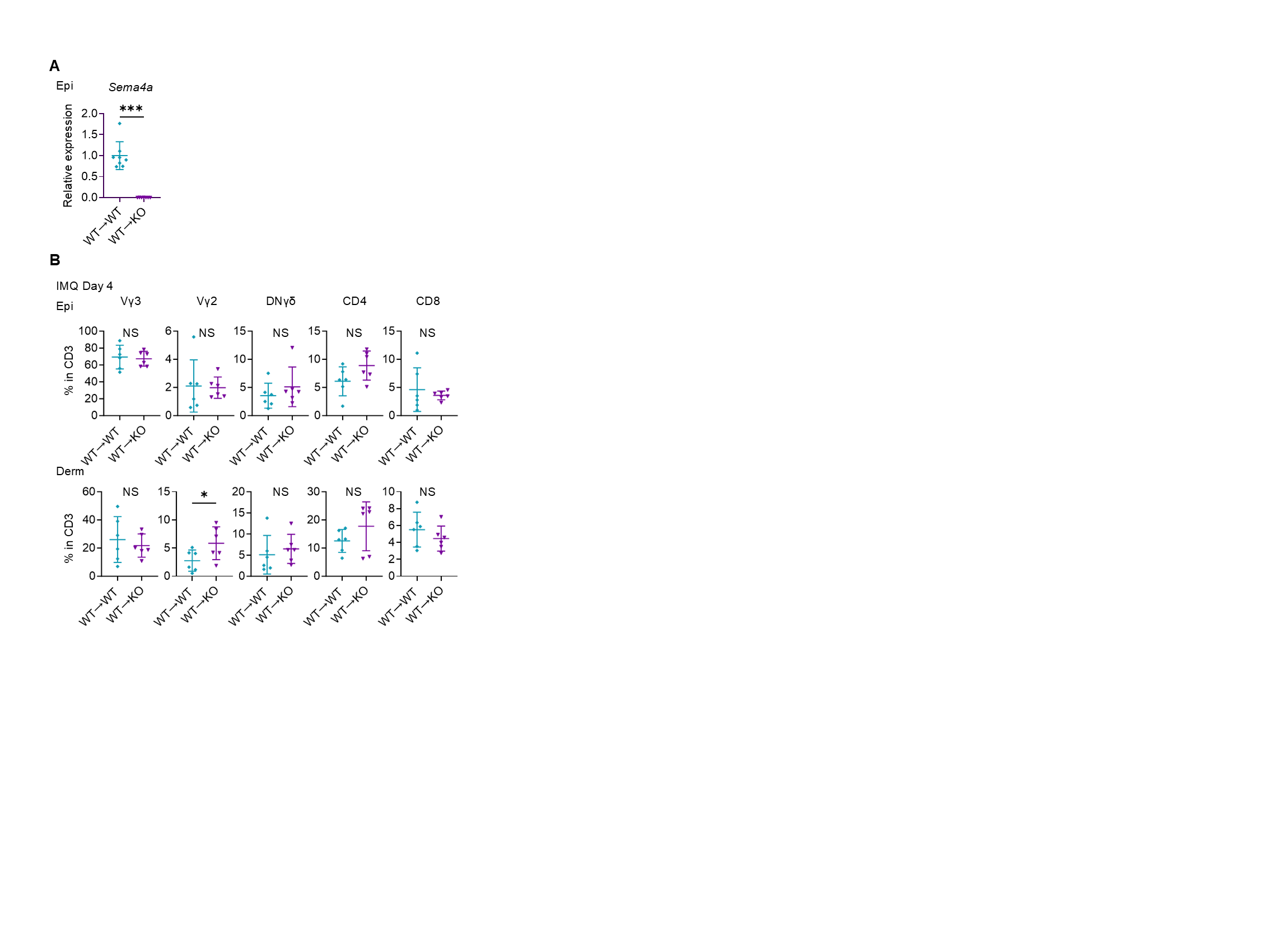

### Figure 4-figure supplement 1

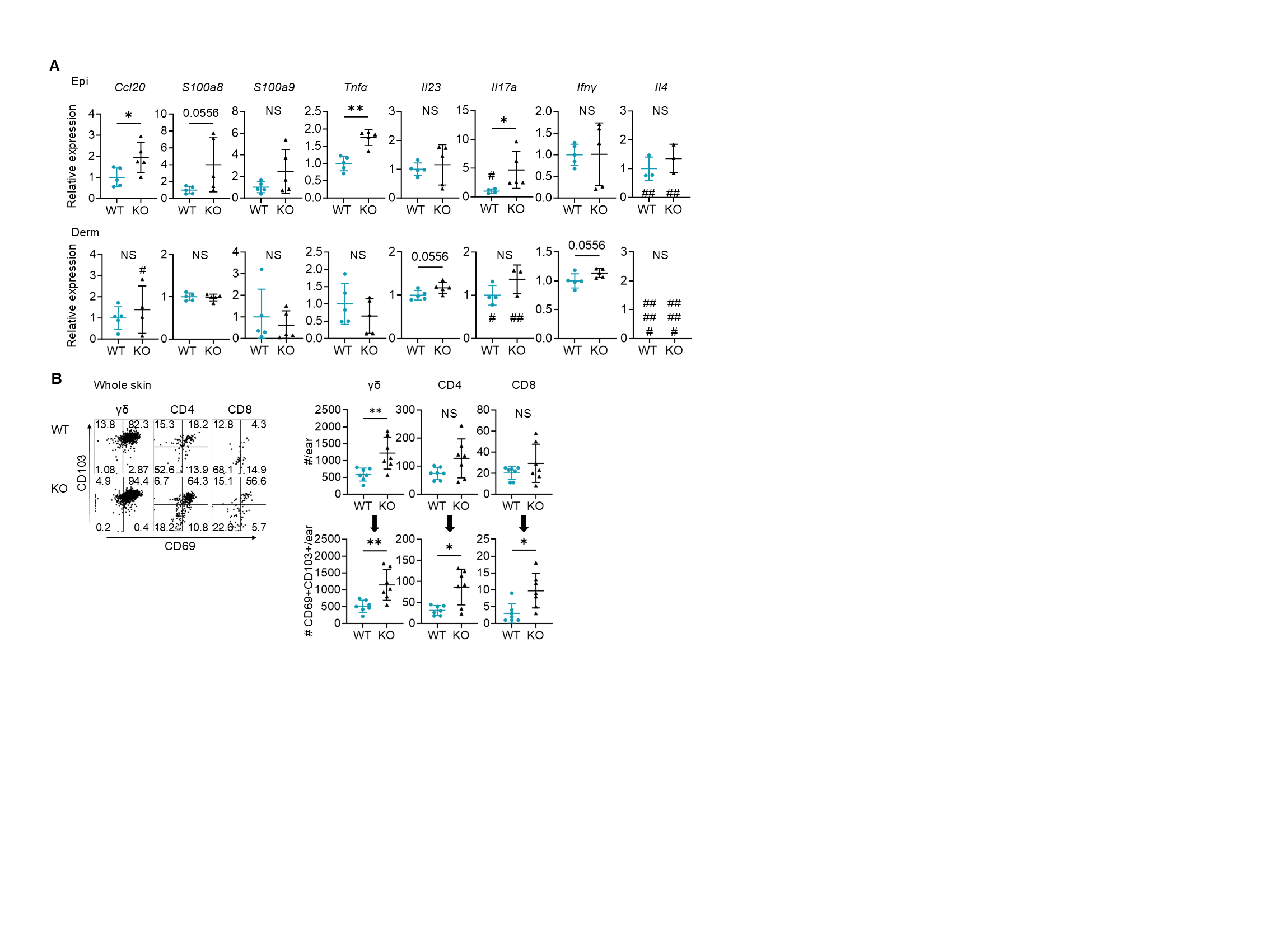

### Figure 4-figure supplement 2

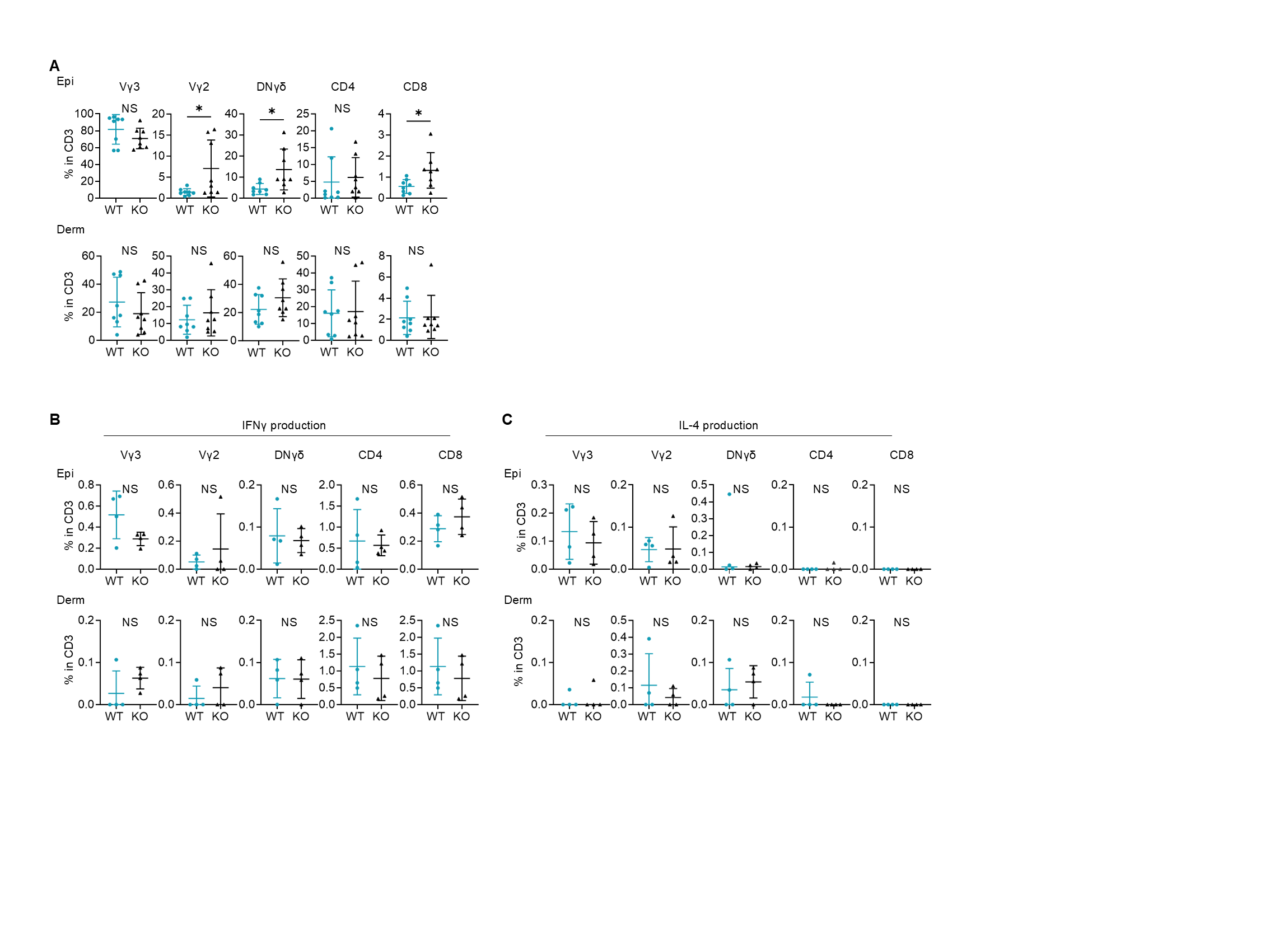

### Figure 4-figure supplement 3

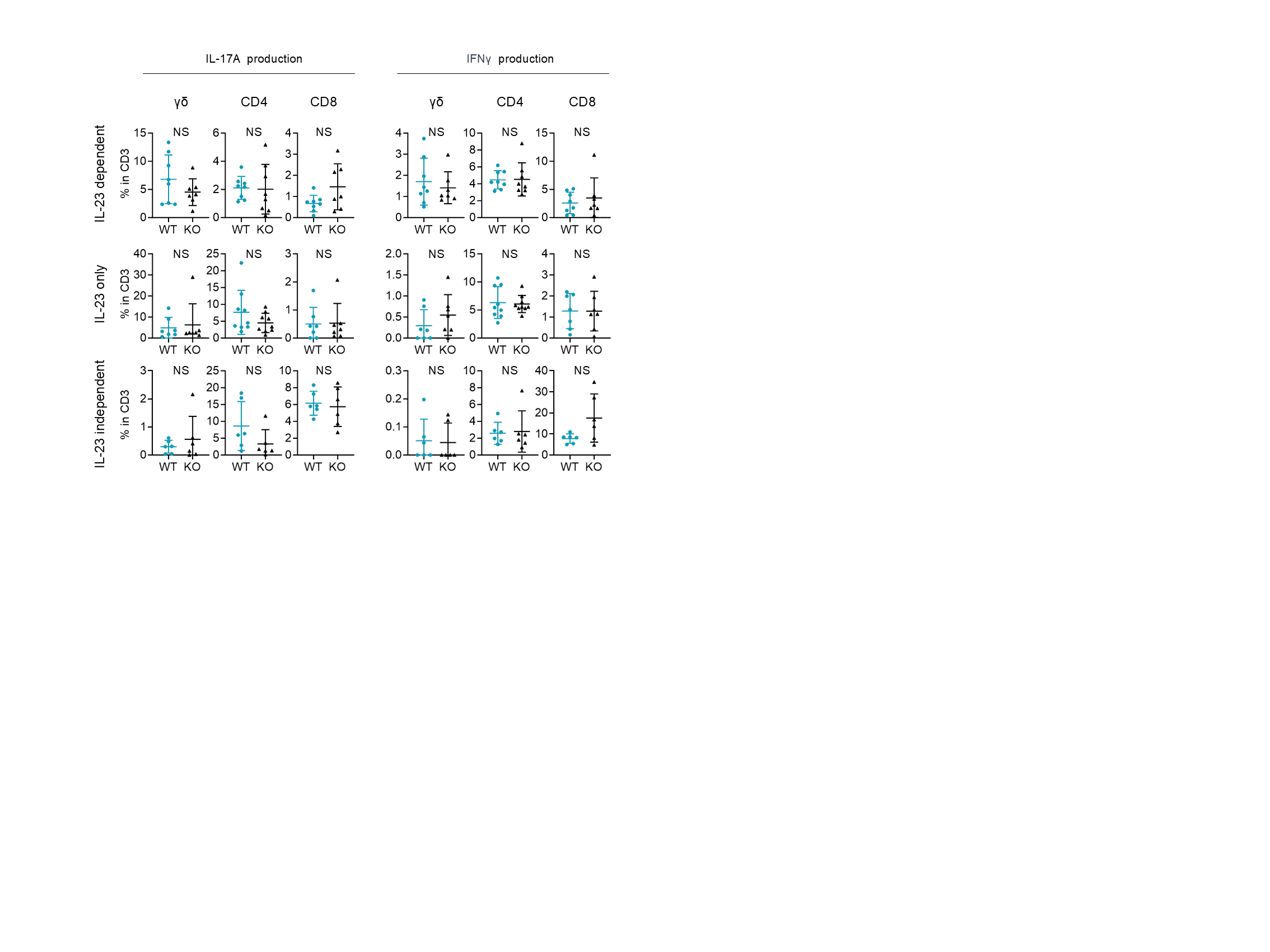

### Figure 5-figure supplement 1

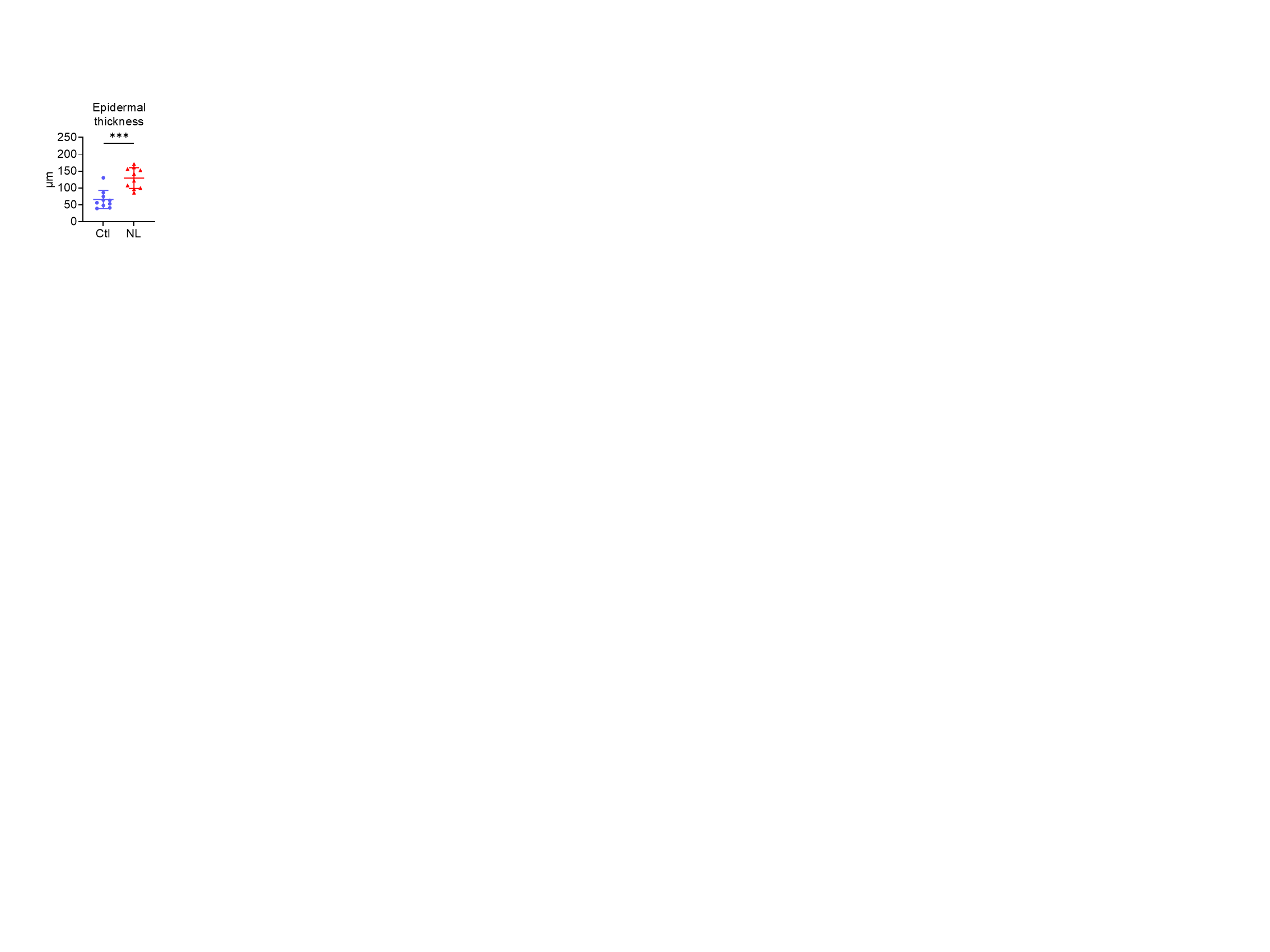

### Figure 5-figure supplement 2

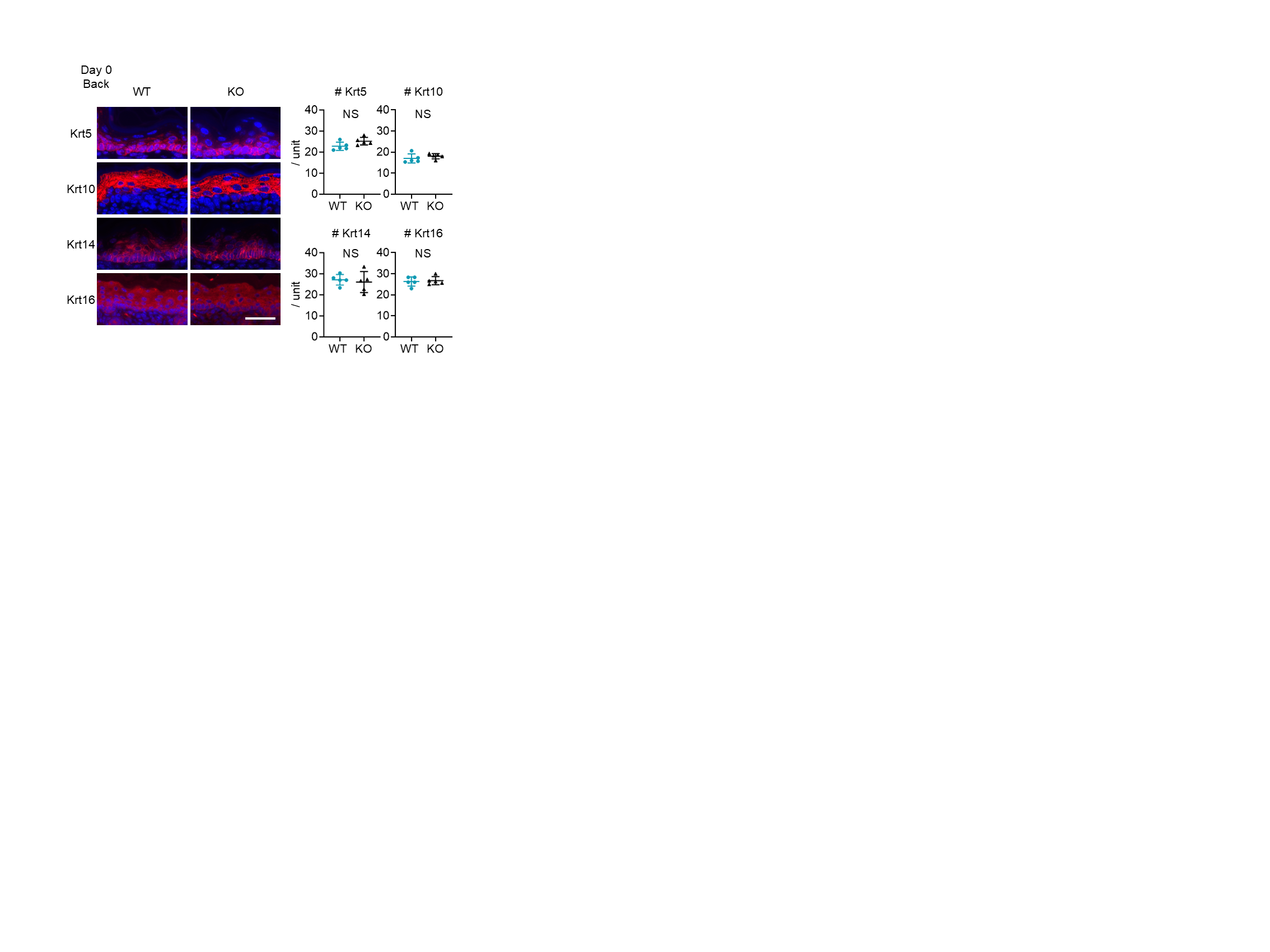

### Figure 7-figure supplement 1

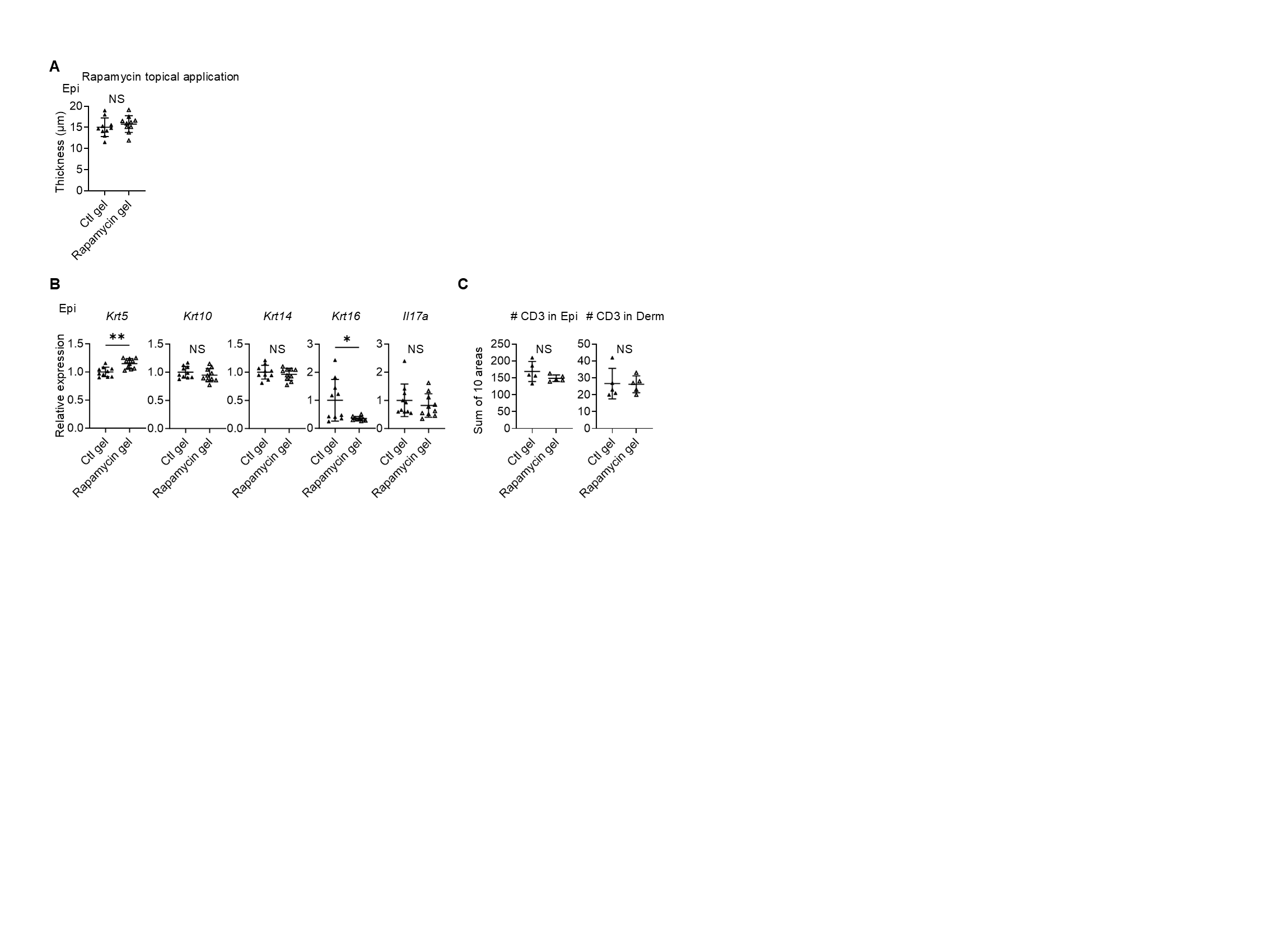

### Figure 7-figure supplement 2

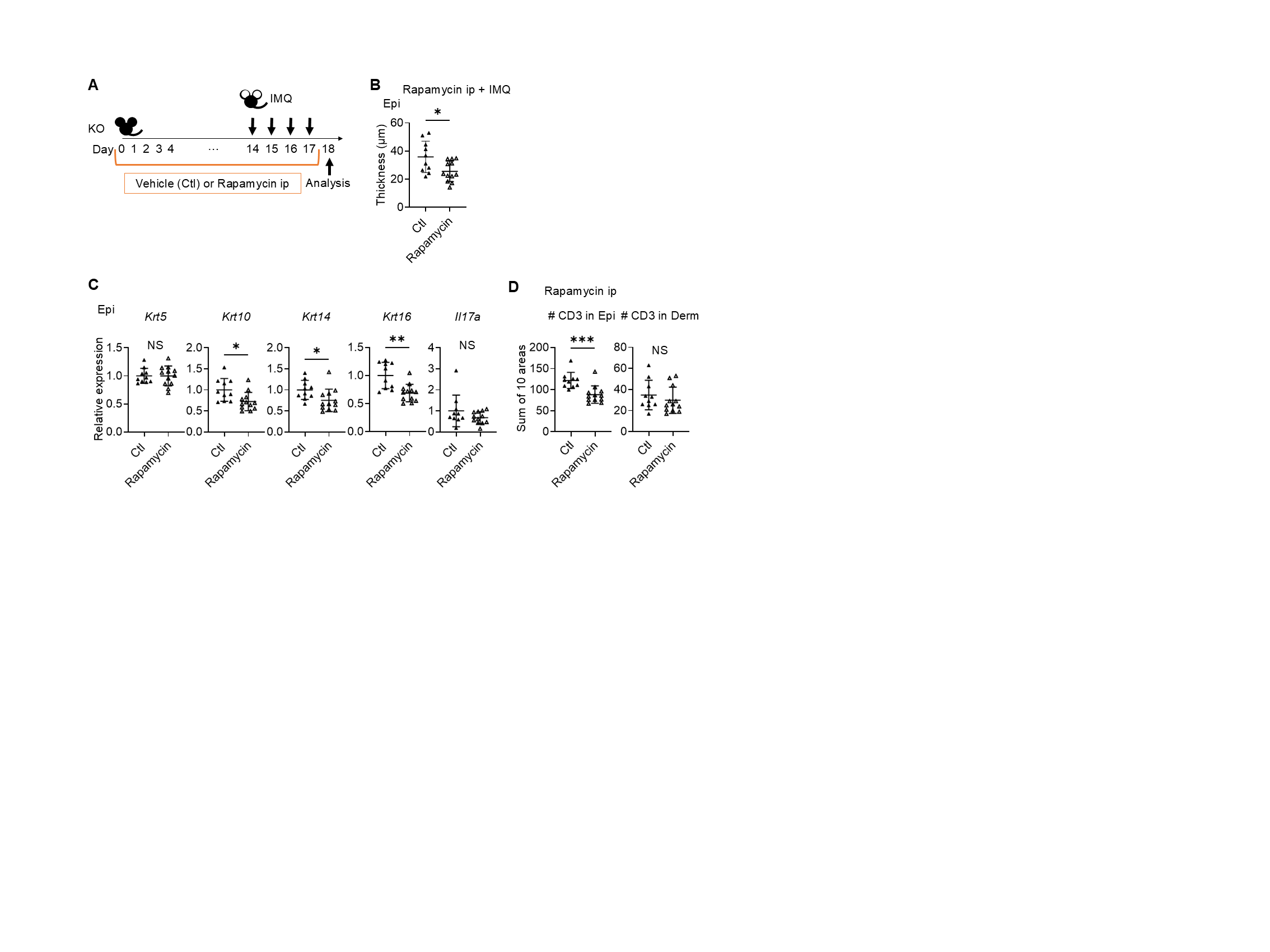
