## Supplemental Tables for "Downregulation of Semaphorin 4A in keratinocytes reflects the features of non-lesional psoriasis"

| Table S1 | Patient information | |  |  |  |  |  |  |
| --- | --- | --- | --- | --- | --- | --- | --- | --- |
|  | Psoriasis severity is defined by the total body surface area (BSA) affected: <3% BSA for mild, 3%-10% BSA for moderate, and >10% BSA for severe disease. | | | | |  |  |  |
| Fig. 1, D and E | Analysis of Sema4A expression by immunohistochemistry | | | |  |  |  |  |
| Fig. S4 | Epidermal thickness | |  |  |  |  |  |  |
|  | Psoriasis (n = 10) | |  |  |  |  |  |  |
|  | Age | Gender | Body site | Severity |  |  |  |  |
|  | 66 | Male | Leg | Severe |  |  |  |  |
|  | 55 | Male | Trunk | Moderate |  |  |  |  |
|  | 80 | Female | Trunk | Severe |  |  |  |  |
|  | 72 | Male | Trunk | Moderate |  |  |  |  |
|  | 30 | Male | Trunk | Severe |  |  |  |  |
|  | 72 | Female | Leg | Moderate |  |  |  |  |
|  | 63 | Male | Arm | Mild |  |  |  |  |
|  | 38 | Male | Trunk | Mild |  |  |  |  |
|  | 35 | Male | Leg | Mild |  |  |  |  |
|  | 71 | Male | Trunk | Moderate |  |  |  |  |
|  | Control (n = 10) | |  |  |  |  |  |  |
|  | Age | Gender | Body site |  |  |  |  |  |
|  | 76 | Male | Neck |  |  |  |  |  |
|  | 52 | Female | Leg |  |  |  |  |  |
|  | 66 | Female | Leg |  |  |  |  |  |
|  | 33 | Male | Leg |  |  |  |  |  |
|  | 68 | Female | Trunk |  |  |  |  |  |
|  | 57 | Female | Leg |  |  |  |  |  |
|  | 65 | Male | Trunk |  |  |  |  |  |
|  | 41 | Female | Trunk |  |  |  |  |  |
|  | 51 | Female | Trunk |  |  |  |  |  |
|  | 66 | Female | Trunk |  |  |  |  |  |
| Fig 1F | Analysis of Sema4A expression by qRT-PCR | | |  |  |  |  |  |
|  | Epidermis |  |  |  |  |  |  |  |
|  | Psoriasis (n = 7) | |  |  |  |  |  |  |
|  | Age | Gender | Body site | Severity |  |  |  |  |
|  | 71 | Male | Leg | Moderate |  |  |  |  |
|  | 76 | Female | Neck | Mild |  |  |  |  |
|  | 38 | Female | Trunk | Moderate |  |  |  |  |
|  | 49 | Female | Trunk | Moderate |  |  |  |  |
|  | 61 | Male | Arm | Mild |  |  |  |  |
|  | 68 | Female | Trunk | Moderate |  |  |  |  |
|  | 72 | Male | Leg | Mild |  |  |  |  |
|  | Control (n = 10) | |  |  |  |  |  |  |
|  | Age | Gender | Body site |  |  |  |  |  |
|  | 57 | Female | Leg |  |  |  |  |  |
|  | 69 | Female | Trunk |  |  |  |  |  |
|  | 66 | Female | Trunk |  |  |  |  |  |
|  | 86 | Female | Trunk |  |  |  |  |  |
|  | 53 | Female | Trunk |  |  |  |  |  |
|  | 84 | Male | Neck |  |  |  |  |  |
|  | 37 | Female | Trunk |  |  |  |  |  |
|  | 46 | Female | Trunk |  |  |  |  |  |
|  | 51 | Female | Trunk |  |  |  |  |  |
|  | 46 | Female | Trunk |  |  |  |  |  |
|  | Dermis |  |  |  |  |  |  |  |
|  | Psoriasis (n = 6) | |  |  |  |  |  |  |
|  | Age | Gender | Body site | Severity |  |  |  |  |
|  | 76 | Female | Neck | Mild |  |  |  |  |
|  | 38 | Female | Trunk | Moderate |  |  |  |  |
|  | 49 | Female | Trunk | Moderate |  |  |  |  |
|  | 61 | Male | Arm | Severe |  |  |  |  |
|  | 68 | Female | Trunk | Moderate |  |  |  |  |
|  | 72 | Male | Leg | Severe |  |  |  |  |
|  | Control (n = 6) | |  |  |  |  |  |  |
|  | Age | Gender | Body site |  |  |  |  |  |
|  | 69 | Female | Trunk |  |  |  |  |  |
|  | 86 | Female | Trunk |  |  |  |  |  |
|  | 84 | Male | Neck |  |  |  |  |  |
|  | 37 | Female | Trunk |  |  |  |  |  |
|  | 51 | Female | Trunk |  |  |  |  |  |
|  | 46 | Female | Trunk |  |  |  |  |  |
| Fig 1G | Analysis of Sema4A expression by flow cytometry | | |  |  |  |  |  |
|  |  | | Cell type | | | Average of age | Male (n = ) | Female (n = ) |
|  | Psoriasis (n = 13) | | CD4 T cell, CD8 T cell, and monocyte | | | 56.1 | 9 | 4 |
|  | Control (n = 13) | | CD4 T cell and CD8 T cell | | | 52.2 | 6 | 7 |
|  | Control (n = 11) | | Monocyte | | | 54.1 | 6 | 5 |
| Fig 1H | Analysis of Serum Sema4A by ELISA | | |  |  |  |  |  |
|  |  | | Average of age | Male (n = ) | Female (n = ) |  |  |  |
|  | Psoriasis (n = 60) | | 55.3 | 43 | 17 |  |  |  |
|  | Control (n = 20) | | 52.1 | 11 | 9 |  |  |  |
| Fig 6A | Pictures of p-S6, S6, p-Akt, and Akt staining | | | | | |  |  |
|  | Psoriasis (n = 1) | |  |  |  |  |  |  |
|  | Age | Gender | Body site | Severity |  |  |  |  |
|  | 30 | Male | Trunk | Severe |  |  |  |  |
|  | Control (n = 1) | |  |  |  |  |  |  |
|  | Age | Gender | Body site |  |  |  |  |  |
|  | 66 | Female | Trunk |  |  |  |  |  |
|  | Analysis of p-S6, S6 expression by immunohistochemistry | | | |  |  |  |  |
|  | Psoriasis (n = 9) | |  |  |  |  |  |  |
|  | Age | Gender | Body site | Severity |  |  |  |  |
|  | 66 | Male | Leg | Severe |  |  |  |  |
|  | 55 | Male | Trunk | Moderate |  |  |  |  |
|  | 80 | Female | Trunk | Severe |  |  |  |  |
|  | 72 | Male | Trunk | Moderate |  |  |  |  |
|  | 30 | Male | Trunk | Severe |  |  |  |  |
|  | 72 | Female | Leg | Moderate |  |  |  |  |
|  | 38 | Male | Trunk | Mild |  |  |  |  |
|  | 35 | Male | Leg | Mild |  |  |  |  |
|  | 71 | Male | Trunk | Moderate |  |  |  |  |
|  | Control (n = 9) | |  |  |  |  |  |  |
|  | Age | Gender | Body site |  |  |  |  |  |
|  | 76 | Male | Neck |  |  |  |  |  |
|  | 66 | Female | Leg |  |  |  |  |  |
|  | 33 | Male | Leg |  |  |  |  |  |
|  | 68 | Female | Trunk |  |  |  |  |  |
|  | 57 | Female | Leg |  |  |  |  |  |
|  | 65 | Male | Trunk |  |  |  |  |  |
|  | 41 | Female | Trunk |  |  |  |  |  |
|  | 51 | Female | Trunk |  |  |  |  |  |
|  | 66 | Female | Trunk |  |  |  |  |  |

| Table S2 |  |  |  |  |  |  |  |  |
| --- | --- | --- | --- | --- | --- | --- | --- | --- |
| Antibodies used for immunohistochemical, immunofluorescence, and western blot analyses | | | | | | | | |
| Target | Clone | Species | Supplier | Catalog | Dilution ratio | | Incubation  time | Incubation  temperature |
|  |  |  |  |  | IHC, IF | WB |  |  |
| Sema4A | Polyclonal | Rabbit | Abcam | ab70178 | 100 |  | 30 min | RT |
| phospho-S6 (Ser235/236) | D57.2.2E | Rabbit | Cell Signaling | 4858 | 400 | 2000 | overnight | 4 °C |
| S6 | 5G10 | Rabbit | Cell Signaling | 2217 | 100 | 1000 | overnight | 4 °C |
| phospho-Akt (Ser473) | D9E | Rabbit | Cell Signaling | 4060 | 50 | 1000 | overnight | 4 °C |
| Akt | C67E7 | Rabbit | Cell Signaling | 4691 | 300 | 1000 | overnight | 4 °C |
| Keratin 5 | Polyclonal | Rabbit | BioLegend | 905503 | 800 |  | overnight | 4 °C |
| Keratin 10 | Polyclonal | Rabbit | BioLegend | 905403 | 400 |  | overnight | 4 °C |
| Keratin 14 | Polyclonal | Rabbit | BioLegend | 905303 | 400 |  | overnight | 4 °C |
| Cytokeratin 16 | 8L6R4 | Rabbit | Invitrogen | MA5-42892 | 100 |  | overnight | 4 °C |
| CD3 | CD3-12 | Rat | Bio-Rad | MCA1477 | 100 |  | overnight | 4 °C |
| Rabbit IgG H&L (Alexa Fluor 555) | Polyclonal | Donkey | Abcam | ab150074 | 1000 |  | 30 min | RT |
| Rat IgG H&L (Alexa Fluor 555) | Polyclonal | Donkey | Abcam | ab150154 | 1000 |  | 30 min | RT |
| β-actin | AC-15 | Mouse | Sigma-Aldrich | A5441 |  | 5000 | overnight | 4 °C |
| Anti-Mouse IgG, HRP-Linked Whole Ab | Monoclonal secondary | Sheep | Cytiva | NA931 |  | 10000 | 1 hour | RT |
| Anti-Rabbit IgG, HRP-Linked Whole Ab | Polyclonal secondary | Donkey | Cytiva | NA934 |  | 10000 | 1 hour | RT |
| RT: Room temperature |  |  |  |  |  |  |  |  |
| IHC: immunohistochemistry |  |  |  |  |  |  |  |  |
| IF: immunofluorescence |  |  |  |  |  |  |  |  |
| WB: western blot |  |  |  |  |  |  |  |  |

| Table S3 |  |  |
| --- | --- | --- |
| Primer sequences of real-time quantitative PCR used in human sample experiments | | |
| Gene Symbol | Forward primer (5'-3') | Reverse primer (5'-3') |
| *GAPDH* | GTCTCCTCTGACTTCAACAGCG | ACCACCCTGTTGCTGTAGCCAA |
| *SEMA4A* | TCTGCTCCTGAGTGGTGATG | AAACCAGGACACGGATGAAG |
| Primer sequences of real-time quantitative PCR in murine sample experiments | | |
| Gene Symbol | Forward primer (5'-3') | Reverse primer (5'-3') |
| *Gapdh* | TGTGTCCGTCGTGGATCTGA | TTGCTGTTGAAGTCGCAGGAG |
| *Krt5* | CAGAGCTGAGGAACATGCAG | CATTCTCAGCCGTGGTACG |
| *Krt10* | CGTACTGTTCAGGGTCTGGAG | GCTTCCAGCGATTGTTTCA |
| *Krt14* | CGTACTGTTCAGGGTCTGGAG | GCTTCCAGCGATTGTTTCA |
| *Krt16* | TGAGCTGACCCTGTCCAGA | TGAGCTGACCCTGTCCAGA |
| *Filaggrin* | GGAGGCATGGTGGAACTGA | TGTTTATCTTTTCCCTCACTTCTACATC |
| *Loricrin* | TCACTCATCTTCCCTGGTGCTT | GTCTTTCCACAACCCACAGGA |
| *Sema4a* | ATGGCCCTACCATCCCTGG | AGCAGCGTGTCAAAGTCTCG |
| *Ccl20* | ATGGCCTGCGGTGGCAAGCGT | CATCTTCTTGACTCTTAGGC |
| *S100a8* | TGCGATGGTGATAAAAGTGG | GGCCAGAAGCTCTGCTACTC |
| *S100a9* | CACCCTGAGCAAGAAGGAAT | TGTCATTTATGAGGGCTTCATTT |
| *Ifnγ* | ATCTGGAGGAACTGGCAAAA | TTCAAGACTTCAAAGAGTCTGAGGTA |
| *Tnfα* | GCCTCCCTCTCATCAGTTCT | CACTTGGTGGTTTGCTACGA |
| *Il4* | ACAGGAGAAGGGACGCCAT | GAAGCCCTACAGACGAGCTCA |
| *Il17a* | CTGTGTCTCTGATGCTGTTG | ATGTGGTGGTCCAGCTTTC |
| *Il23* | TCCCTACTAGGACTCAGCCAAC | GCTGCCACTGCTGACTAGAA |

| Table S4 |  |  |  |  |
| --- | --- | --- | --- | --- |
| Antibodies used for flow cytometry analysis | | | |  |
| Target | Clone | Species | Supplier | Catalog |
| CD3 | OKT3 | Mouse | BioLegend | 317335 |
| CD4 | RPA-T4 | Mouse | BioLegend | 300512 |
| CD8a | RPA-T8 | Mouse | eBioscience | 47-0088-42 |
| CD11c | 3.9 | Mouse | BioLegend | 301628 |
| SEMA4A | 5E3/SEMA4A | Mouse | BioLegend | 148404 |
| CD3ε | 145-2C11 | Armenian Hamster | BioLegend | 100328 |
| CD4 | GK1.5 | Rat | BioLegend | 100406 |
| CD8a | 53-6.7 | Rat | BioLegend | 100714 |
| CD16/32 | 93 | Rat | BioLegend | 101301 |
| CD69 | H1.2F3 | Armenian Hamster | BioLegend | 104514 |
| CD103 | 2E7 | Armenian Hamster | BioLegend | 121422 |
| TCR Vγ2 | UC3-10A6 | Armenian Hamster | BioLegend | 137705 |
| TCR Vγ3 | 536 | Syrian Hamster | BD Biosciences | 743241 |
| TCRγδ | GL3 | Armenian Hamster | BioLegend | 118124 |
| IFNγ | XMG1.2 | Rat | BioLegend | 505813 |
| IL-4 | 11B11 | Rat | BD Biosciences | 562915 |
| IL-17A | [TC11-18H10.1](https://www.biolegend.com/ja-jp/search-results?Clone=TC11-18H10.1) | Rat | BioLegend | 506925 |

| Table S5 | |  |
| --- | --- | --- |
| Mouse recombinant cytokines | | |
| Cytokines | Supplier | Catalog |
| IL-1β | BioLegend | 575102 |
| IL-6 | BioLegend | 575702 |
| IL-23 | BioLegend | 589002 |
| TGFβ1 | BioLegend | 763102 |
