## Supplementary material for "Downregulation of Semaphorin 4A in keratinocytes reflects the features of non-lesional psoriasis": Key Resources Table

| **Key Resources Table** | | | | |
| --- | --- | --- | --- | --- |
| **Reagent type (species) or resource** | **Designation** | **Source or reference** | **Identifiers** | **Additional information** |
| strain, strain background (*Mus musculus*) | C57BL/6J WT mice | CLEA Japan |  |  |
| strain, strain background (*Mus musculus*) | Semaphorin 4A knockout mice | Dr. Atsushi Kumanogoh (Osaka University, Osaka, Japan) |  |  |
| biological sample (Homo sapiens) | Skin specimens from 17 psoriasis patients | Osaka University |  |  |
| biological sample (Homo sapiens) | Skin specimens from 19 subjects who underwent tumor resection or reconstructive surgery | Osaka University |  |  |
| biological sample (Homo sapiens) | Blood samples from 73 psoriasis patients | Osaka University |  |  |
| biological sample (Homo sapiens) | Blood samples from 33 Ctl | Osaka University |  |  |
| antibody | Sema4A (Rabbit polyclonal) | Abcam | Cat# Ab70178; RRID: AB_1270611 | IHC (1:100) |
| antibody | phospho-S6 (Ser235/236) (Rabbit monoclonal) | Cell Signaling Technology | Cat# 4858 | IHC (1:400), WB (1:2000) |
| antibody | S6 (Rabbit monoclonal) | Cell Signaling Technology | Cat# 2217 | IHC (1:100), WB (1:1000) |
| antibody | phospho-Akt (Ser473) (Rabbit monoclonal) | Cell Signaling Technology | Cat# 4060 | IHC (1:50), WB (1:1000) |
| antibody | Akt (Rabbit monoclonal) | Cell Signaling Technology | Cat# 4691 | IHC (1:300), WB (1:1000) |
| antibody | Keratin 5 (Rabbit polyclonal) | BioLegend | Cat# 905503; RRID: AB_2734679 | IF (1:800) |
| antibody | Keratin 10 (Rabbit polyclonal) | BioLegend | Cat# 905403; RRID: AB_2749902 | IF (1:400) |
| antibody | Keratin 14 (Rabbit polyclonal) | BioLegend | Cat# 905303; RRID: AB_2734678 | IF (1:400) |
| antibody | Cytokeratin 16 (Rabbit monoclonal) | Invitrogen | Cat# MA5-42892; RRID: AB_2912033 | IF (1:100) |
| antibody | CD3 (Rat monoclonal) | Bio-Rad | Cat# MCA1477; RRID: AB_321245 | IF (1:100) |
| antibody | Rabbit IgG H&L (Alexa Fluor 555) (Donkey polyclonal) | Abcam | Cat# ab150074; RRID: AB_2636997 | IF (1:1000) |
| antibody | Rat IgG H&L (Alexa Fluor 555) (Donkey polyclonal) | Abcam | Cat# ab150154; RRID: AB_2813834 | IF (1:1000) |
| antibody | β-actin (Mouse monoclonal) | Sigma-Aldrich | Cat# A5441 | WB (1:5000) |
| antibody | Anti-Mouse IgG, HRP-Linked Whole Ab (Sheep monoclonal secondary) | Cytiva | Cat# NA931; RRID: AB_772210 | WB (1:10000) |
| antibody | Anti-Rabbit IgG, HRP-Linked Whole Ab (Donkey polyclonal secondary) | Cytiva | Cat# NA934; RRID: AB_772206 | WB (1:10000) |
| antibody | CD3 (Mouse monoclonal) | BioLegend | Cat# 317335; RRID: AB_2561627 | FCM (1:100) |
| antibody | CD4 (Mouse monoclonal) | BioLegend | Cat# 300512; RRID: AB_314080 | FCM (1:100) |
| antibody | CD8a (Mouse monoclonal) | eBioscience | Cat# 47-0088-42; RRID:  AB_1272046 | FCM (1:100) |
| antibody | CD11c (Mouse monoclonal) | BioLegend | Cat# 301628; RRID: AB_11203895 | FCM (1:100) |
| antibody | SEMA4A (Mouse monoclonal) | BioLegend | Cat# 148404; RRID: AB_2565287 | FCM (1:100) |
| antibody | CD3ε (Armenian Hamster monoclonal) | BioLegend | Cat# 100328; RRID: AB_893318 | FCM Skin specimens (1:20), Others (1:100) |
| antibody | CD4 (Rat monoclonal) | BioLegend | Cat# 100406; RRID: AB_312691 | FCM (1:100) |
| antibody | CD8a (Rat monoclonal) | BioLegend | Cat# 100714; RRID: AB_312753 | FCM (1:100) |
| antibody | CD16/32 (Rat monoclonal) | BioLegend | Cat# 101301; RRID: AB_312800 | FCM (1:100) |
| antibody | CD69 (Armenian Hamster monoclonal) | BioLegend | Cat# 104514; RRID: AB_492843 | FCM (1:10) |
| antibody | CD103 (Armenian Hamster monoclonal) | BioLegend | Cat# 121422; RRID: AB_2562901 | FCM (1:100) |
| antibody | TCR Vγ2 (Armenian Hamster monoclonal) | BioLegend | Cat# 137705; RRID:  AB_10643997 | FCM (1:100) |
| antibody | TCR Vγ3 (Syrian Hamster monoclonal) | BD Biosciences | Cat# 743241; RRID: AB_2741371 | FCM (1:100) |
| antibody | TCRγδ (Armenian Hamster monoclonal) | BioLegend | Cat# 118124; RRID: AB_11204423 | FCM (1:100) |
| antibody | IFNγ (Rat monoclonal) | BioLegend | Cat# 505813; RRID: AB_493312 | FCM (1:40) |
| antibody | IL-4 (Rat monoclonal) | BD Biosciences | Cat# 562915; RRID: AB_2737889 | FCM (1:40) |
| antibody | IL-17A (Rat monoclonal) | BioLegend | Cat# 506925; RRID: AB_10900442 | FCM (1:40) |
| sequence-based reagent | human *GAPDH*_F | This paper | PCR primers | GTCTCCTCTGACTTCAACAGCG |
| sequence-based reagent | human *GAPDH*_R | This paper | PCR primers | ACCACCCTGTTGCTGTAGCCAA |
| sequence-based reagent | human *SEMA4A*_F | (Carvalheiro et al., 2019) | PCR primers | TCTGCTCCTGAGTGGTGATG |
| sequence-based reagent | human *SEMA4A*_R | (Carvalheiro et al., 2019) | PCR primers | AAACCAGGACACGGATGAAG |
| peptide, recombinant protein | Recombinant Mouse IL-1β (carrier-free) | BioLegend | Cat# 575102 |  |
| peptide, recombinant protein | Recombinant Mouse IL-6 (carrier-free) | BioLegend | Cat# 575702 |  |
| peptide, recombinant protein | Recombinant Mouse IL-23 (carrier-free) | BioLegend | Cat# 589006 |  |
| peptide, recombinant protein | Recombinant Mouse TGF-β1 (carrier-free) | BioLegend | Cat# 763102 |  |
| commercial assay or kit | BD Cytofix/Cytoperm^TM^ Fixation/Permeabilization Kit | BD Biosciences | Cat# 554714 |  |
| commercial assay or kit | Dako REAL EnVision Detection System, Peroxidase/DAB, Rabbit/Mouse, HRP kit | Agilent | Cat# K5007; RRID: AB_2888627 |  |
| commercial assay or kit | Direct-zol RNA Miniprep Kits | ZYMO Research | Cat# R2050 |  |
| commercial assay or kit | High-Capacity RNA-to-cDNA™ Kit | Thermo Fisher scientific | Cat# 4387406 |  |
| commercial assay or kit | LIVE/DEAD^TM^ Fixable Dead Cell Stain Kit | Thermo Fisher Scientific | Cat# L34965 |  |
| commercial assay or kit | Pan T Cell Isolation Kit II, mouse | Miltenyi Biotec | Cat# 130-095-130 |  |
| commercial assay or kit | T Cell Activation/Expansion Kit, mouse | Miltenyi Biotec | Cat# 130-093-627 |  |
| commercial assay or kit | TB Green Premix Ex Taq^TM^ II (Tli RNaseH Plus) | Takara-bio | Cat# RR820A |  |
| chemical compound, drug | 5％ imiquimod cream | Mochida | Global Trade Item Number: 224130002 |  |
| chemical compound, drug | BD Golgiplug^TM^ | BD Biosciences | Cat# 555029 |  |
| chemical compound, drug | Collagenase type III | Worthington Biochemical Corporation | Cat# LS004183 |  |
| chemical compound, drug | CountBright™ Absolute Counting Beads, for flow cytometry | Thermo Fisher Scientific | Cat# C36950 |  |
| chemical compound, drug | Dispase Ⅱ | Wako | Cat# 383-02281 |  |
| chemical compound, drug | Ionomycin | WAKO | Cat# 095-05831 |  |
| chemical compound, drug | JR-AB2-011 | MedChemExpress | Cat# HY-122022 |  |
| chemical compound, drug | Mounting medium with DAPI | Vector Laboratories | Cat# H-1200; RRID:　AB_2336790 |  |
| chemical compound, drug | Phorbol 12-Myristate 13-Acetate | WAKO | Cat# 162-23591 |  |
| chemical compound, drug | Phosphatase Inhibitor Cocktail(100x) | Nacalai tesque | Cat# 07574-61 |  |
| chemical compound, drug | Protease Inhibitor Cocktail for Use with Mammalian Cell and Tissue Extracts | Nacalai tesque | Cat# 25955-11 |  |
| chemical compound, drug | Protein Block Serum-Free | Agilent | Cat# X0909 |  |
| chemical compound, drug | Rapamycin | Sanxin Chempharma | CAS# 53123-88-9 |  |
| chemical compound, drug | RBC Lysis Buffer (10X) | BioLegend | Cat# 420301 |  |
| chemical compound, drug | WB Stripping Solution | Nacalai tesque | Cat# 05364-55 |  |
| software, algorithm | Cellxgene VIP | DOI: https://doi.org/10.1101/2020.08.28.270652 | (K. Li et al., 2022) |  |
| software, algorithm | GraphPad Prism 10 | GraphPad Software | RRID: SCR_002798 |  |
| software, algorithm | Image J | National Institutes of Health | RRID:SCR_003070 |  |
| software, algorithm | Kaluza | Beckman Coulter | RRID: SCR_016182 |  |
| software, algorithm | RaNAseq | https://ranaseq.eu/ | (Prieto & Barrios, 2019) |  |
